# Cell-Hub: a graphical interface for end-to-end single-cell RNA sequencing analysis

**DOI:** 10.64898/2026.07.15.738621

**Authors:** Gaspard Macaux, Maxime Di Gallo, Valentina Taglietti, Helge Amthor, Pascal Maire

**Author notes:** Corresponding author: Gaspard Macaux, 24 rue du Faubourg Saint-Jacques, 75014 Paris, France.

## Abstract

Single-cell and single-nucleus RNA sequencing have become increasingly widespread, creating a significant demand for accessible analysis tools in research laboratories. Despite this need, the bioinformatics expertise required for such analyses remains rare. Cell-Hub addresses this gap by enabling single-cell data analysis for all researchers, regardless of computational background.

Cell-Hub is a comprehensive, free, and open-source framework built on R/Shiny and distributed as a Docker image, integrating Seurat 5, CellChat 2, and Monocle 3 within a unified graphical interface. It supports all essential steps of single-cell RNA-seq analysis: data loading, quality control, normalization, clustering, multi-dataset integration, differential expression, and biomarker detection. Cell-Hub further incorporates ligand-receptor interaction inference powered by GaspouDB, a consolidated database of 11,563 mouse and 9,604 human interactions derived from CellChat, CellPhoneDB, CellTalkDB, and MultiNicheNet as well as trajectory inference via Monocle 3 and spatial transcriptomics analysis for 10X Visium datasets. All analyses produce publication-ready visualizations with flexible export options.

By integrating these analytical frameworks into a single, intuitive interface requiring no programming expertise, Cell-Hub represents a significant step toward democratizing single- cell genomics for the broader research community.

## Introduction

Single-cell RNA sequencing (scRNA-seq) technologies have revolutionized our understanding of cellular heterogeneity by enabling the profiling of gene expression at unprecedented resolution. However, the high dimensionality and complexity of scRNA-seq datasets often present substantial analytical barriers, as many biological research groups lack the specialized computational expertise needed to effectively process and interpret these data without dedicated bioinformatics support.

To address this gap, we present Cell-Hub, a comprehensive and user-friendly framework designed to democratize single-cell data analysis. Built using R and the Shiny framework, Cell- Hub provides a graphical interface that abstracts the complexity of code-based workflows while maintaining scientific rigor. The platform is distributed as a Docker image for reproducible deployment and integrates Seurat 5^2^ pipelines, enabling researchers to perform end-to-end analysis from quality control and multi-dataset integration to clustering, differential expression, and biomarker detection. Cell-Hub further extends beyond standard transcriptomic profiling to support trajectory inference via Monocle 3^3^ and ligand-receptor interaction analysis via CellChat^4^ (Fig. 1).

**Figure 1.**
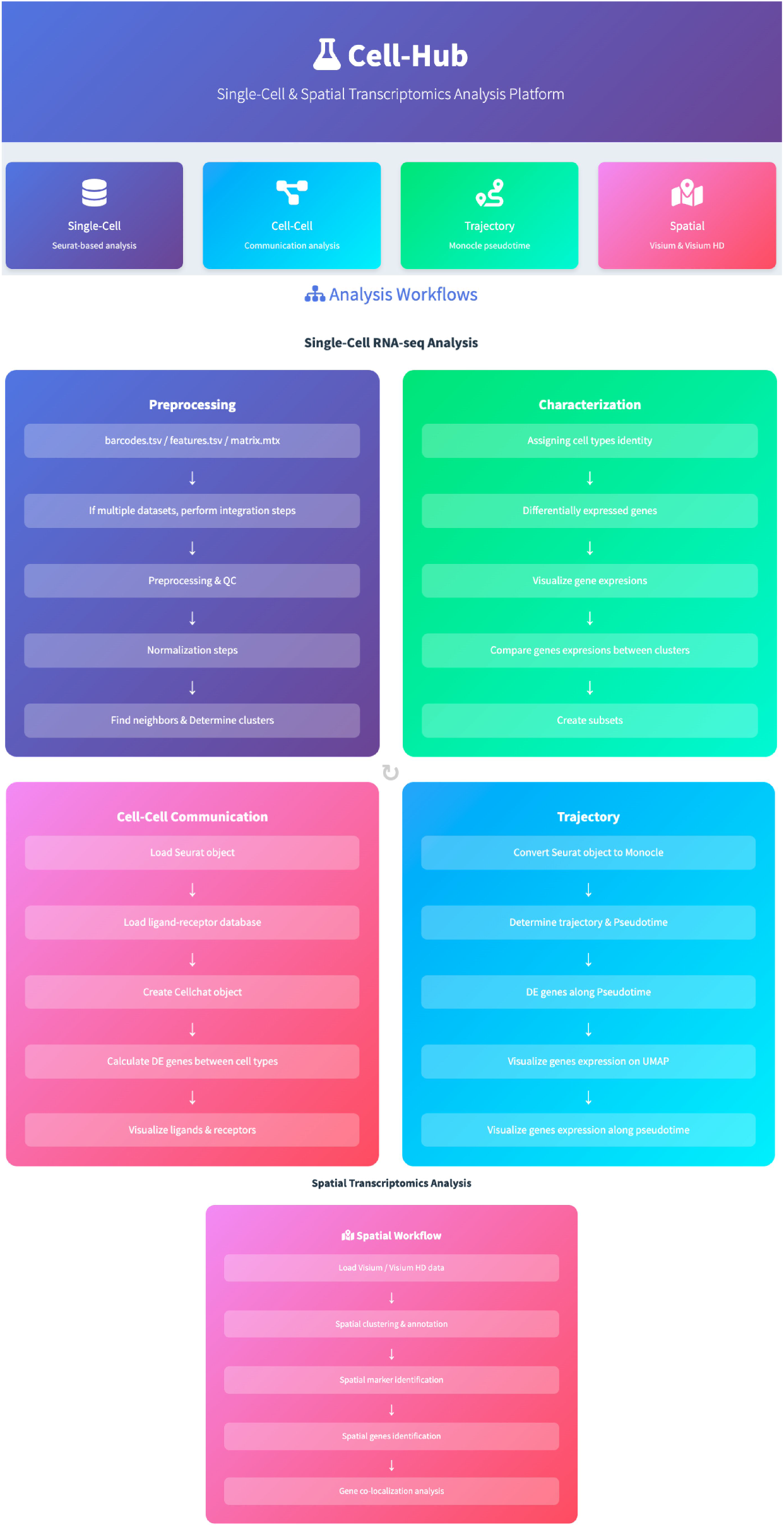
Overview of Cell-Hub analysis workflows and modules. Cell-Hub is a comprehensive web-based platform for single-cell and spatial transcriptomics analysis. The platform integrates four main analytical modules: (1) Single-Cell RNA-seq Analysis, comprising preprocessing steps (data loading, integration, quality control, normalization, and clustering) and characterization workflows (cell type annotation, differential gene expression, visualization, cluster comparison, and subset creation); (2) Cell-Cell Communication Analysis, enabling ligand-receptor interaction inference from Seurat objects using curated databases; (3) Trajectory Analysis, supporting pseudotime inference and differential expression along developmental trajectories via Monocle conversion; and (4) Spatial Transcriptomics Analysis, facilitating spatial clustering, marker identification, and gene localization analysis from Visium and Visium HD datasets.

### Deployment

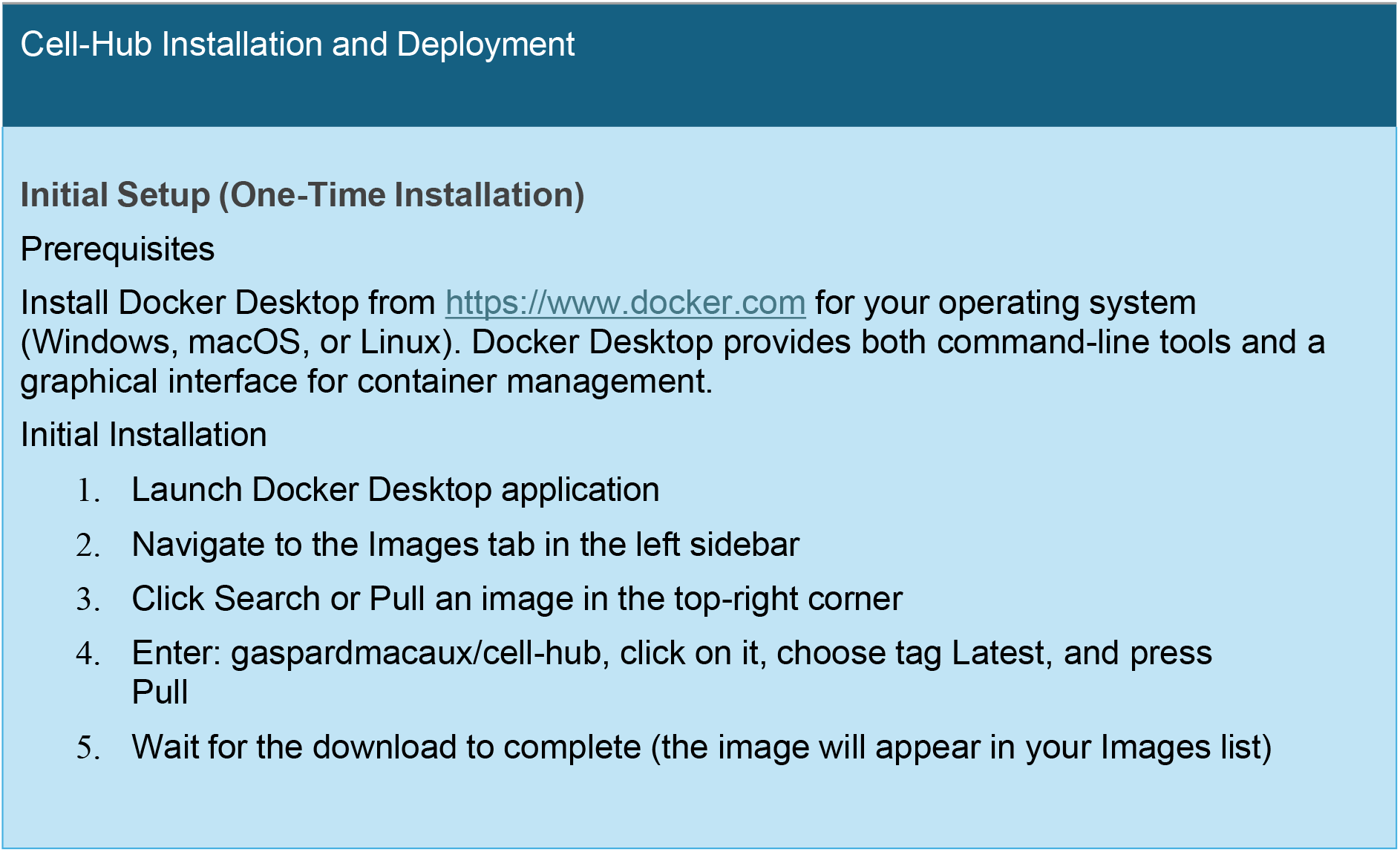

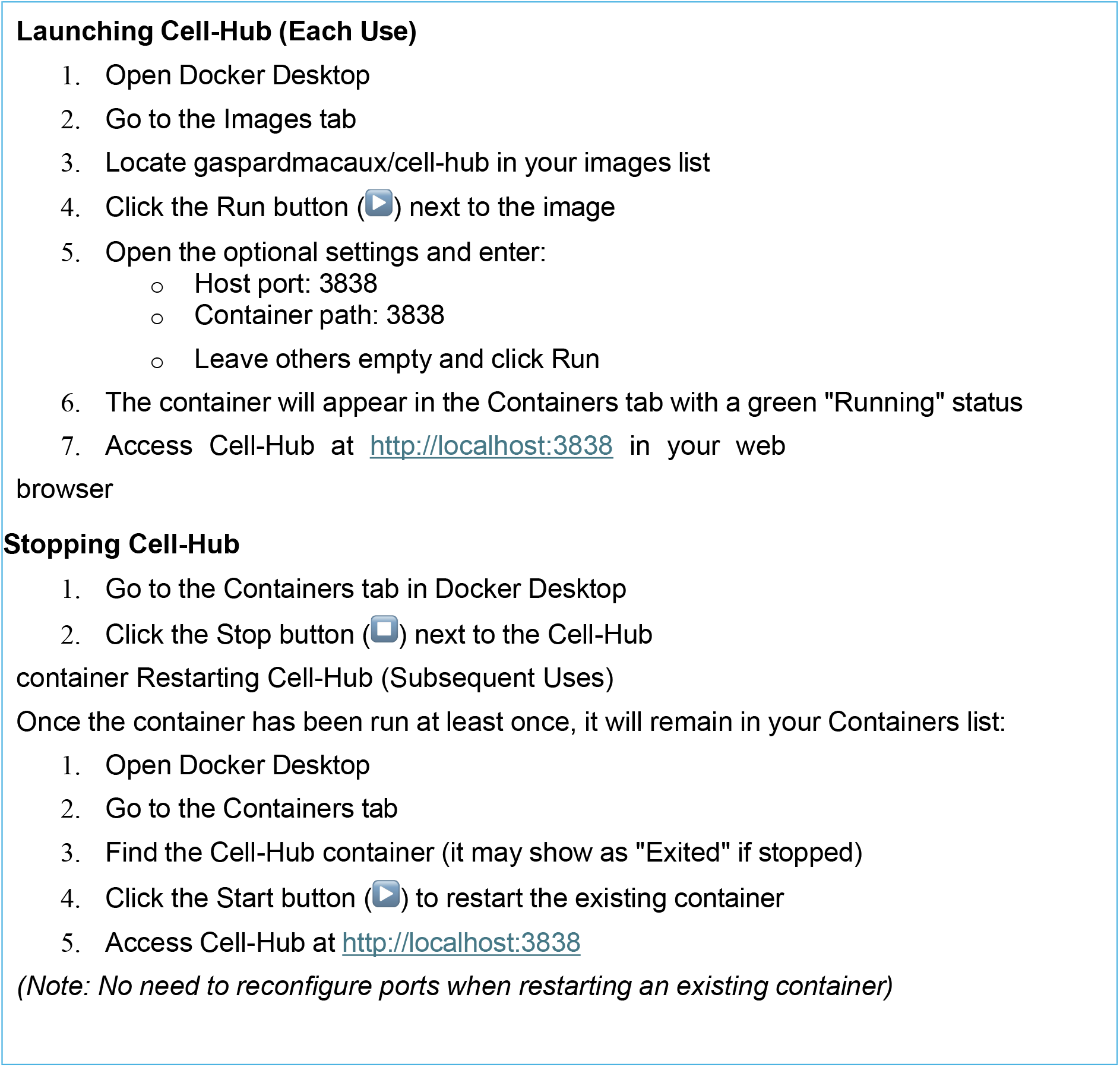

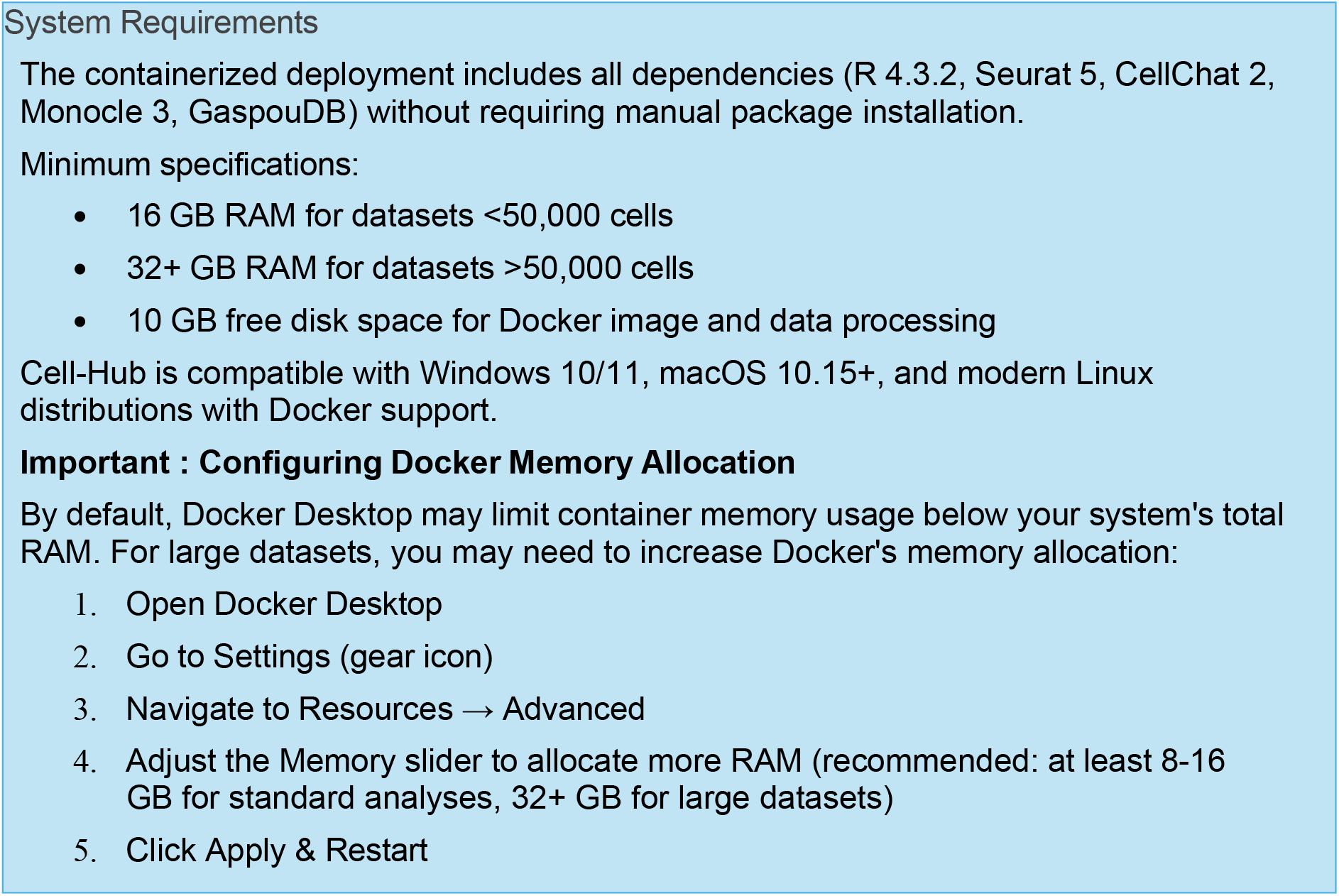

For detailed explanations of analytical parameters and their selection, we direct the reader to the official documentation of the underlying libraries:

– Seurat (https://satijalab.org/seurat/)
– CellChat (https://github.com/jinworks/CellChat)
– Monocle 3 (https://cole-trapnell-lab.github.io/monocle3/).

### Single-Cell Data Analysis with Cell-Hub

#### Data Initialization and Import Workflow

Cell-Hub streamlines the initialization of single-cell workflows through a flexible data import module (Fig. 2). Users first define the experimental context via the Dataset Parameters interface, selecting the target species (*Homo sapiens*, *Mus musculus*) and assay type (Fig. 2a).

**Figure 2.**
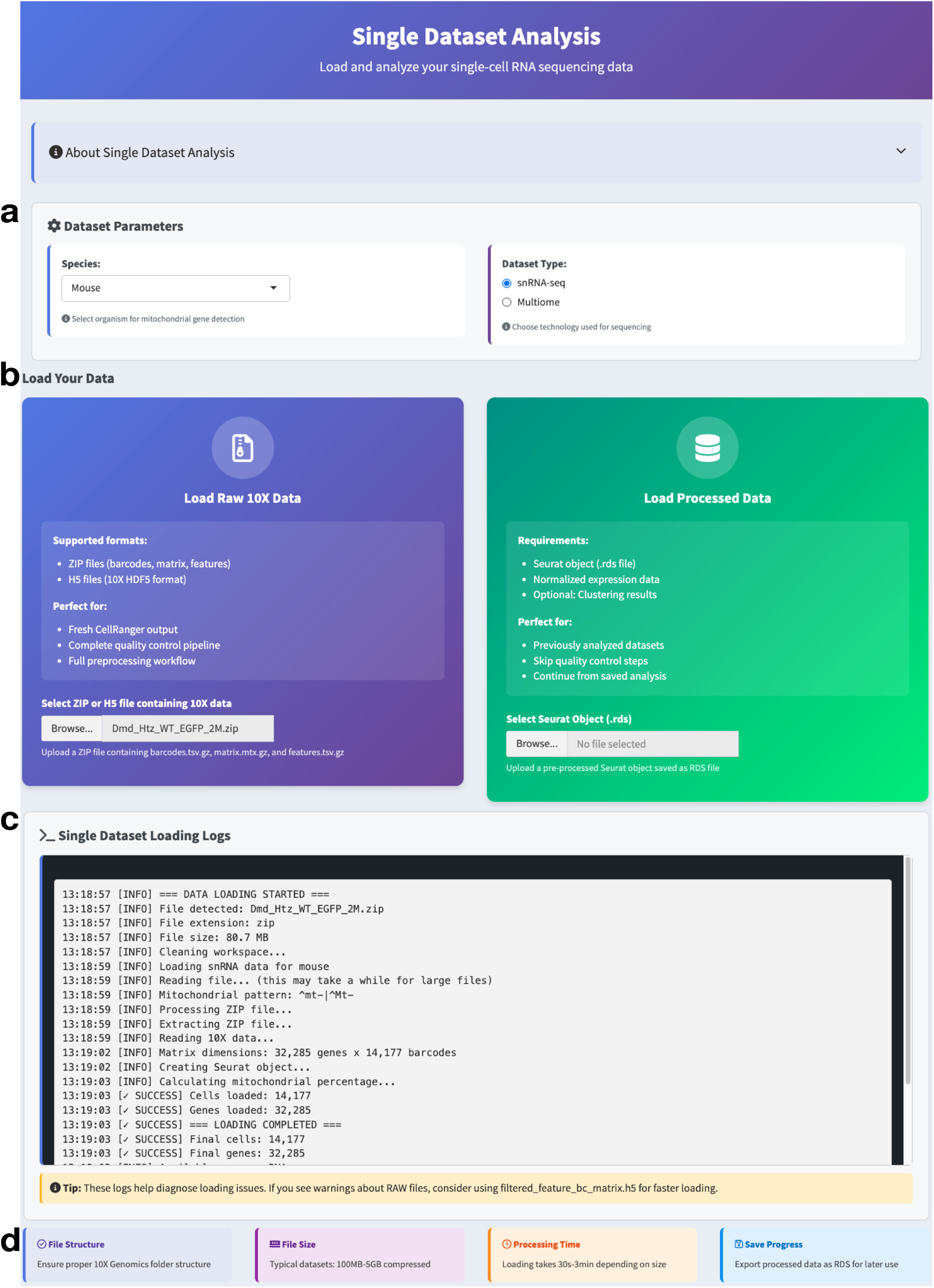
Single dataset analysis interface and data loading workflow. The interface provides two data loading pathways with real-time feedback, (a) Dataset parameter selection for organism and sequencing technology, (b) Data loading options: raw 10X data (left) for complete preprocessing workflow, or pre-processed Seurat objects (right) to continue from previous analyses, (c) Real-time console logs tracking data loading progress and validation, (d) Information panels displaying file structure requirements, typical file sizes, processing time estimates, and data export options.

The Data Loading module supports widely used input formats: raw 10X Genomics outputs (barcodes, features, and matrix files compressed as a ZIP archive), 10X HDF5 (.h5) and AnnData (.h5ad) files, and pre-processed Seurat objects (.rds). Files are uploaded via the integrated graphical upload interface (Fig. 2b), eliminating the need for manual directory structuring or command-line file handling. Upon upload, the system automatically initializes the Seurat object and ensures correct formatting for downstream analysis.

A real-time Log Console reports the status of background processes, confirming successful file reading and object creation, and surfacing key dataset metrics (Fig. 2c). To assist researchers unfamiliar with single-cell workflows, the interface includes integrated Tips and Guidelines offering contextual help and formatting instructions to prevent common import errors (Fig. 2d).

#### Quality Control and Normalization

Cell-Hub implements comprehensive quality control (QC) workflows to ensure data integrity before downstream analysis. The QC module provides interactive parameter adjustment (Fig. 3a), allowing users to define filtering thresholds for key metrics including minimum genes per cell, maximum mitochondrial genome percentage, and normalization scale factor. Real-time summary statistics display the total number of cells/nuclei, unique genes detected across the dataset, and median genes per cell/nucleus, providing immediate feedback on filtering stringency.

**Figure 3.**
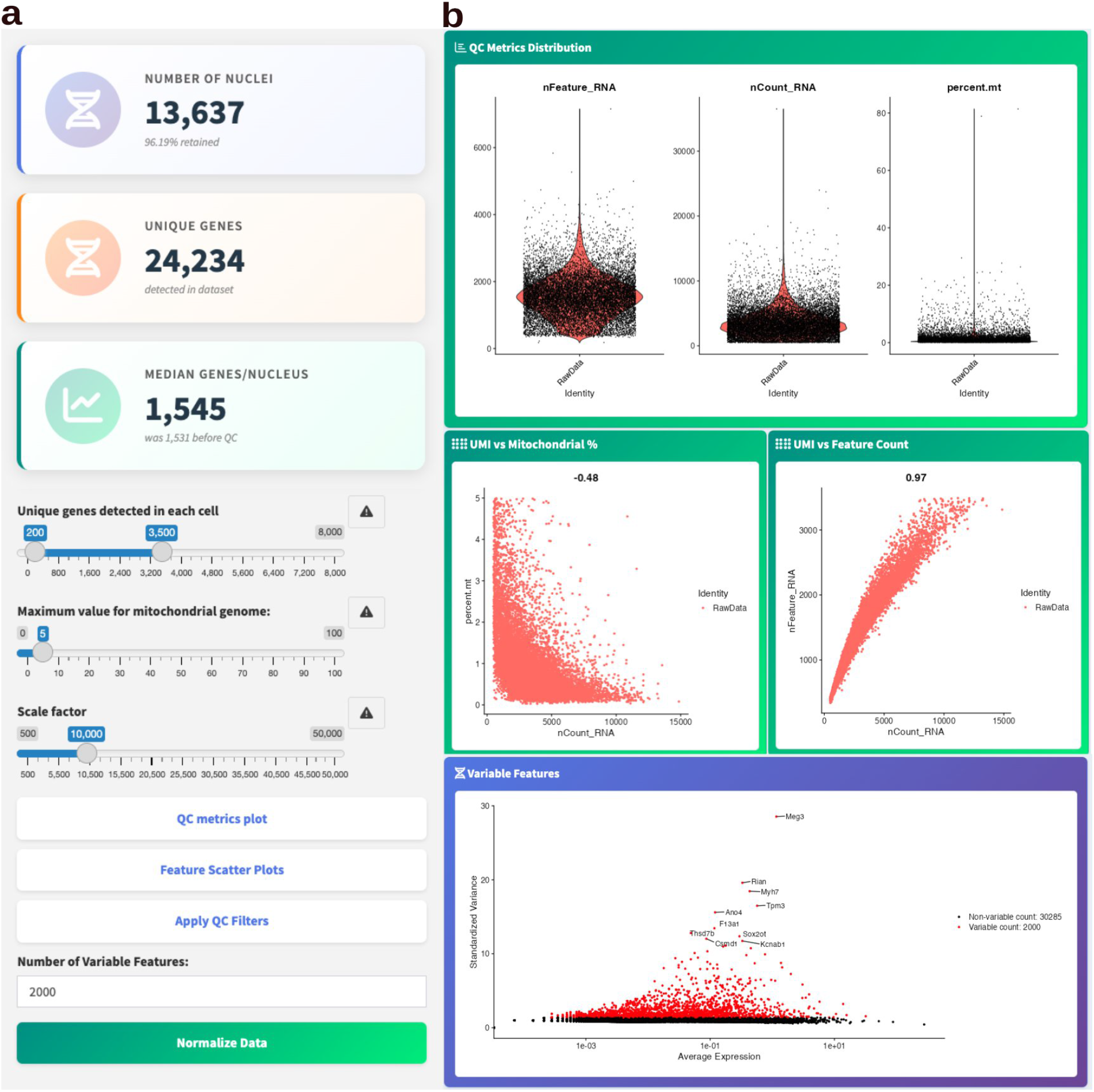
Quality control metrics and filtering interface. (a) Interactive parameter controls for QC filtering, including adjustable thresholds for genes per cell, mitochondrial genome percentage, and normalization scale factor. Summary statistics display total nuclei count, unique genes detected, and median genes per nucleus, (b) QC visualization panels: violin plots for distribution of RNA features, UMI counts, and mitochondrial percentage across cells (top); scatter plots showing relationships between UMI/mitochondrial content and UMI/ feature counts with correlation coefficients (middle); variable features plot identifying highly variable genes for downstream analysis (bottom).

The QC visualization panel (Fig. 3b) presents multiple diagnostic plots to guide filtering decisions. Violin plots display the distribution of RNA features (nFeature_RNA), UMI counts (nCount_RNA), and mitochondrial percentage across all cells/nuclei. Scatter plots examine pairwise relationships between technical metrics: UMI counts versus mitochondrial content, and UMI counts versus feature counts. Cells with high mitochondrial content relative to UMI counts are indicative of stressed or dying cells with compromised membrane integrity, and are flagged for removal alongside potential doublets.

Following quality filtering, Cell-Hub provides four normalization methods adapted to different analytical contexts. LogNormalize performs library-size scaling followed by log-transformation (default scale factor: 10,000); highly variable genes are subsequently identified using variance- stabilizing transformation and displayed as standardized variance against average expression (Fig. 3c). By default, the top 2,000 most variable genes are selected for dimensional reduction. SCTransform performs normalization, variance stabilization, and regression of technical confounders in a single integrated step, without requiring separate highly variable gene identification. CLR (Centered Log-Ratio normalization) is designed for multimodal experiments such as CITE-seq, normalizing each feature across cells to account for the compositional structure of combined RNA and protein measurements. RC (Relative Counts) performs simple library-size scaling without log-transformation, retained for compatibility with downstream tools that require non-log-transformed input.

#### Data Scaling and Principal Component Analysis

Following normalization and feature selection, data are prepared for dimensionality reduction. For LogNormalize and CLR workflows, Cell-Hub scales gene expression values to zero mean and unit variance, preventing highly expressed genes from dominating principal component analysis (PCA). For SCTransform workflows, variance stabilization is performed internally during normalization, eliminating the need for separate scaling.

PCA is executed on the processed data to capture the major axes of transcriptional variation (Fig. 4a). For each PC, the top positive and negative gene loadings are displayed (Fig. 4b), and loading plots visualize the distribution of gene contributions along the PC_1 and PC_2 axes (Fig. 4c), revealing genes with the strongest influence on primary sources of variation.

**Figure 4.**
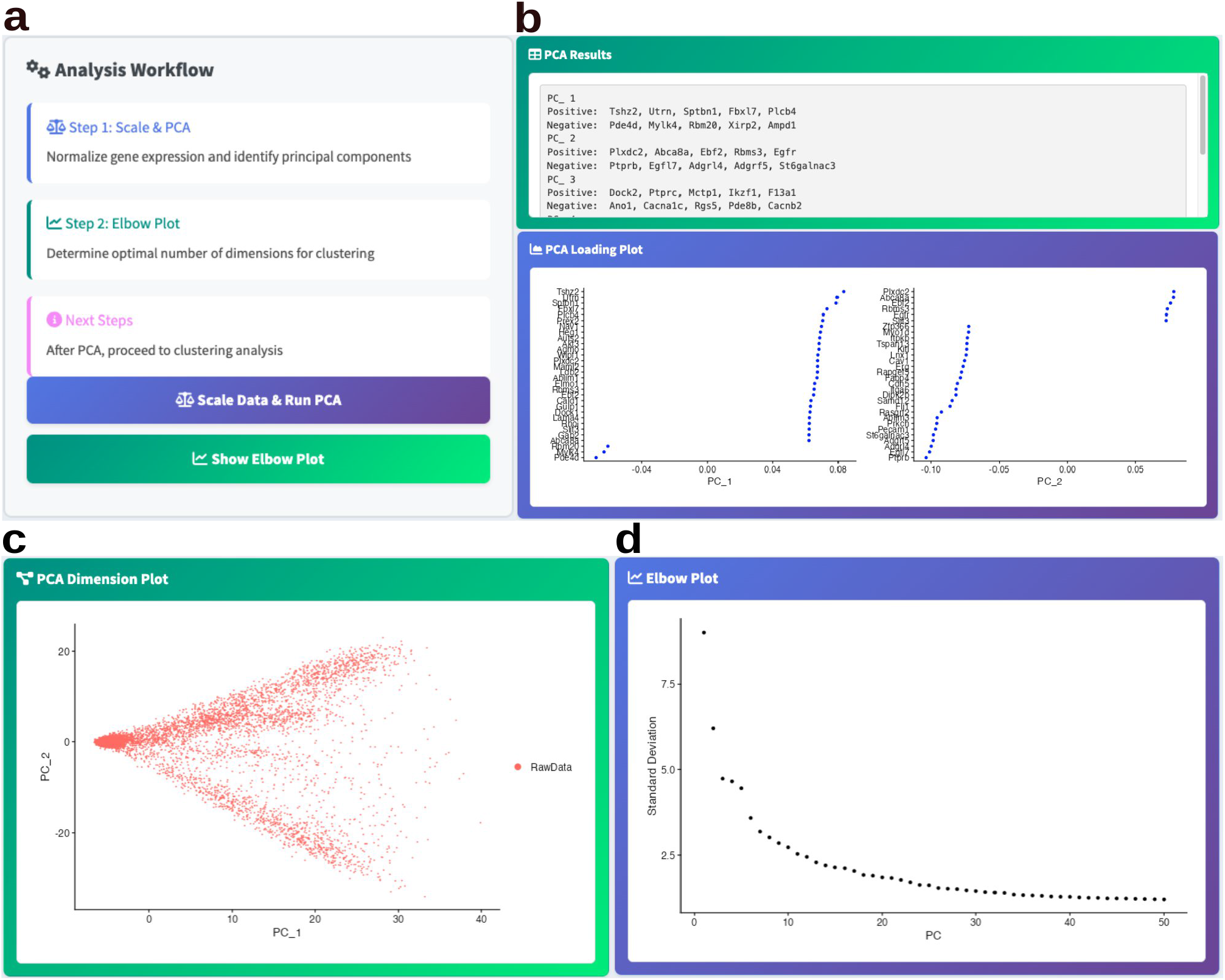
Principal component analysis and dimensionality assessment. **(a)** Control panel with action buttons to execute data scaling and PCA computation, and to generate elbow plot for dimension selection, (b) PCA results displaying top positive and negative gene contributors for each principal component (top), and loading plots showing gene contributions along PC_1 and PC_2 axes (bottom), **(c)** Cell distribution across the first two principal components, **(d)** Elbow plot displaying standard deviation across principal components to identify optimal number of dimensions for downstream clustering.

The PCA dimension plot projects all cells onto the first two principal components (Fig. 4d). To guide selection of the number of informative PCs for downstream clustering, Cell-Hub generates an elbow plot displaying the standard deviation explained by each component (Fig. 4e). The inflection point at which the curve begins to plateau indicates the transition from biologically informative to noise-dominated components, and serves as a practical criterion for PC selection.

#### Cell Clustering

Cell-Hub implements graph-based clustering using a shared nearest neighbor (SNN) approach to identify discrete cell populations. Users configure clustering parameters through an interactive interface (Fig. 5a), including the number of principal components used for neighborhood graph construction (default: 15), the resolution parameter controlling cluster granularity (default: 0.5), and the clustering algorithm. Three algorithms are available: Louvain^8^, Louvain with multilevel refinement^9^, and Smart Local Moving (SLM)^10^. The resolution parameter directly controls cluster number, with higher values producing finer-grained partitions and lower values yielding broader groupings.

**Figure 5.**
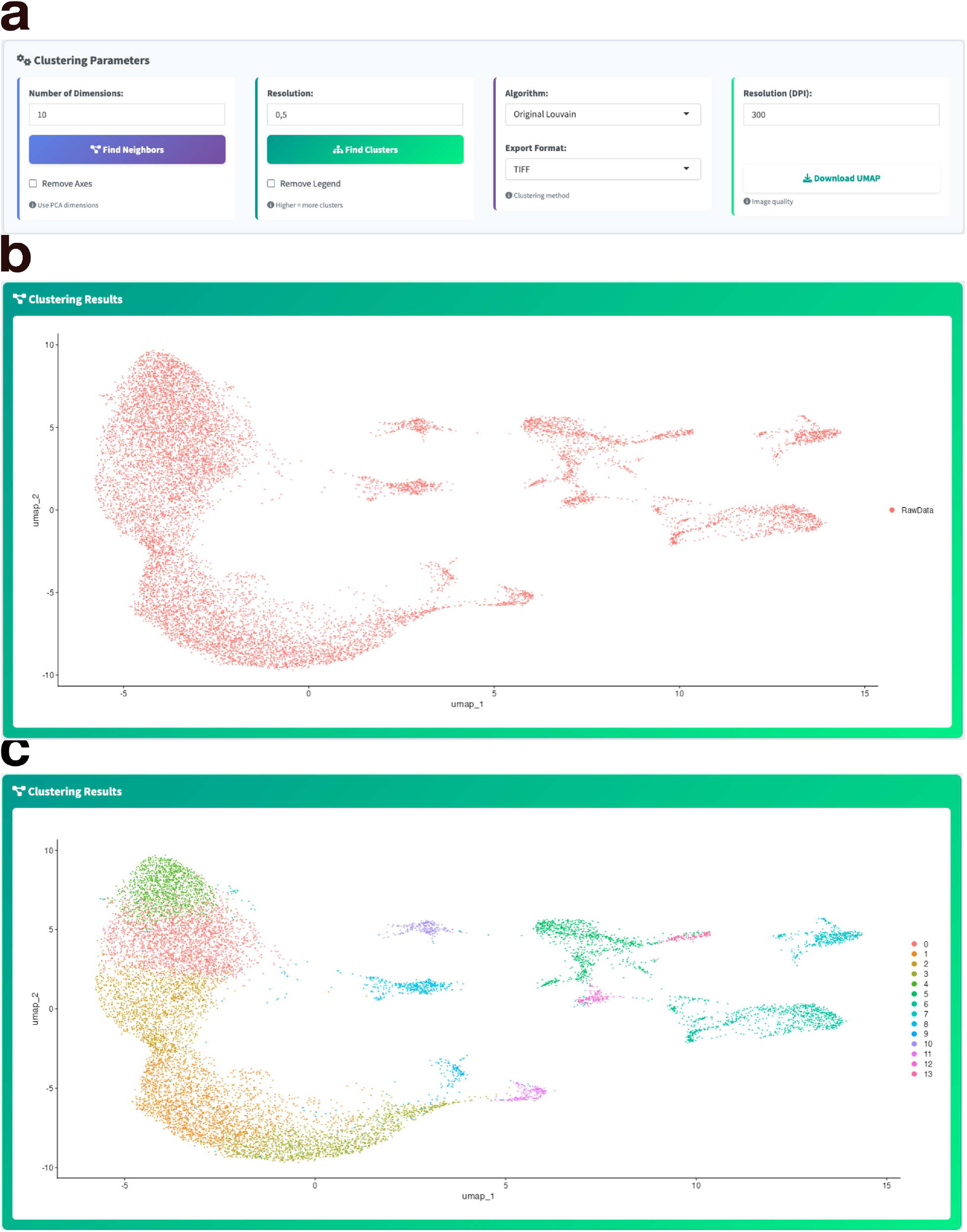
Cell clustering and UMAP visualization. **(a)** Interactive parameter controls for neighborhood graph construction and cluster identification, including adjustable dimensions, resolution, clustering algorithm selection, and UMAP export options with customizable format and resolution, **(b)** UMAP projection displaying overall cell distribution in two-dimensional space, **(c)** UMAP visualization with identified cell clusters color-coded by cluster identity.

Following graph construction, cells are partitioned based on transcriptional similarity using the selected algorithm. To visualize clustering results, Cell-Hub generates Uniform Manifold Approximation and Projection (UMAP) embeddings, projecting cells into two-dimensional space while preserving local neighborhood structure. Both unclustered UMAP projections (Fig. 5b) and cluster-annotated visualizations with color-coded identities (Fig. 5c) are displayed.

To ensure reproducibility, all stochastic steps, including neighborhood graph construction and UMAP embedding, are executed with a fixed random seed (seed = 42), guaranteeing identical results across repeated analyses. UMAP visualizations can be exported in multiple formats (TIFF, PNG, PDF) at adjustable resolution (default: 300 DPI).

#### Doublet Detection and Removal

Cell-Hub implements DoubletFinder^11^ to identify doublets (artificial transcriptomic profiles resulting from co-encapsulation of multiple cells during droplet-based capture). DoubletFinder generates synthetic doublets by averaging the expression profiles of randomly selected cell pairs, then identifies real doublets based on their proximity to these synthetic references in PCA space.

Users configure four key parameters (Fig. 6a): expected doublet rate (default: 7.5%, adjustable according to cell loading density and manufacturer guidelines), number of PCs (default: 10), pN (the proportion of artificial doublets to generate, default: 0.25), and pK (the PC neighborhood size used for doublet scoring, default: 0.09; dataset-specific optimization via parameter sweep is supported). The UMAP projection displays detected doublets in red and singlets in gray (Fig. 6b). Detection statistics (Fig. 6c) report cell counts and percentages before doublet removal, ensuring transparent quality control decisions.

**Figure 6.**
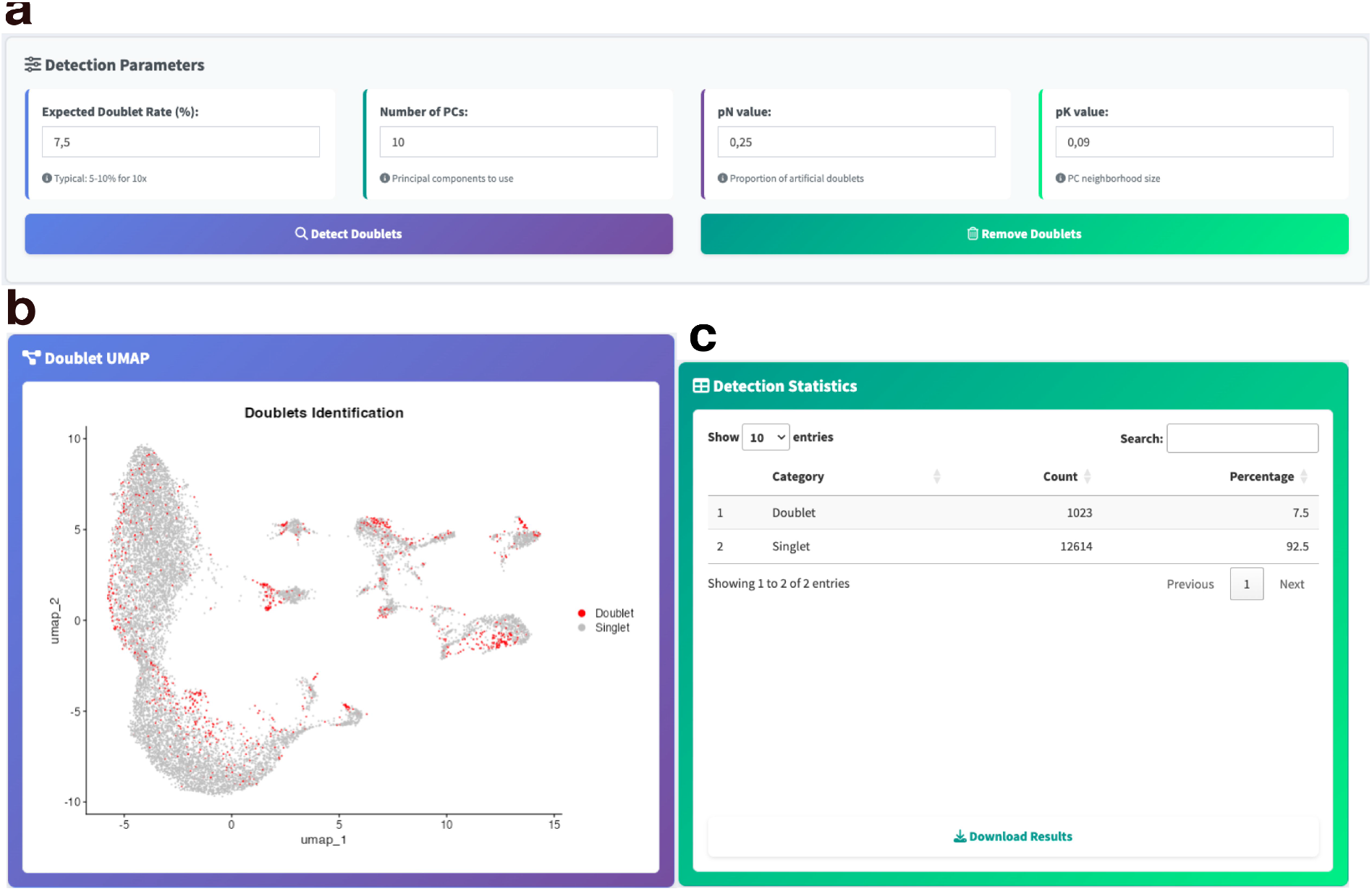
Doublet detection and removal. (a) Parameter controls for doublet detection including expected doublet rate, number of principal components, pN value (proportion of artificial doublets), and pK value (PC neighborhood size), with action buttons to detect and remove doublets, (b) UMAP visualization distinguishing detected doublets (red) from singlets (gray), (c) Detection statistics table displaying counts and percentages of identified doublets and singlets, with download option for results.

#### Gene Expression Visualization

Cell-Hub provides multiple complementary visualization methods to examine gene expression patterns across identified cell populations. A unified parameter panel (Fig. 7a) enables selection of genes of interest, assay type (RNA for log-normalized data, SCT for variance- stabilized data, or integrated for dimensionality-reduction-based multi-sample analyses), export format, and display options applicable across all visualization modules.

**Figure 7.**
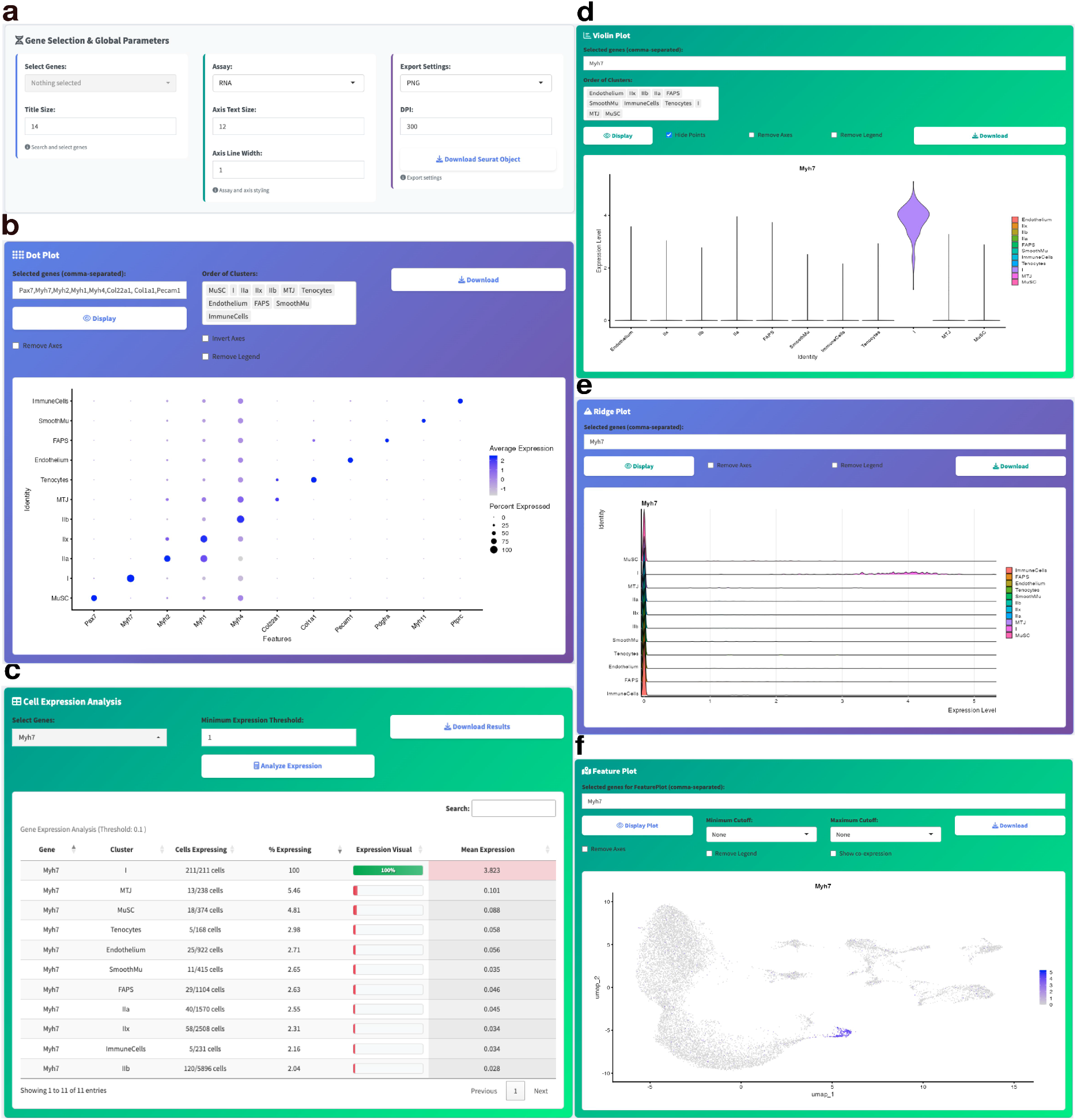
Overview of integrated gene expression visualization modules. (a) Global parameter selection panel enabling customizable gene, assay, and export settings for all visualization modules, (b) Dot plot illustrating gene expression patterns across cell clusters, with dot size corresponding to the percentage of expressing cells and color intensity representing average expression levels, (c) Interactive cell expression analysis table summarizing, for each cluster, the number of cells expressing the selected gene, percentage expressing, and mean expression value, (d) Violin plot depicting the distribution of gene expression levels within each cluster, (e) Ridge plot visualizing the density of gene expression across clusters, (f) Feature plot displaying the spatial distribution of gene expression in two-dimensional UMAP space. Each module supports dynamic selection of analytical parameters and export options for reproducible, user-driven analysis.

Dot plots (Fig. 7b) summarize expression patterns across clusters, with dual encoding: dot size indicates the percentage of expressing cells, while color intensity represents mean expression level. In the multi-dataset module, dot plots can additionally be split by dataset of origin or by any metadata condition. Marker genes with both high percentage expression and high mean expression in a target cluster with minimal off-target signal are readily identifiable from this representation. The expression analysis table (Fig. 7c) provides quantitative metrics including cell counts, expression percentages, and mean values per cluster, exportable in CSV format.

Distribution-based visualizations include violin plots (Fig. 7d), displaying expression distributions within each cluster, and ridge plots (Fig. 7e), providing overlapping density profiles suited for comparing expression patterns across many clusters simultaneously. These complementary approaches facilitate identification of genes with cluster-specific expression patterns.

Feature plots (Fig. 7f) project gene expression onto UMAP space as a continuous color gradient, revealing the spatial distribution of expression across the embedding. Continuous expression gradients across the UMAP may illustrate underlying biological processes such as differentiation trajectories or functional state transitions within a population.

#### Heatmap and Gene Co-expression Analysis

Cell-Hub provides complementary tools for examining gene expression patterns and combinatorial relationships across cell populations. The heatmap module (Fig. 8a) visualizes average expression levels of selected genes across clusters using Z-score normalization per gene, calculated as z = (x - μ)/σ, where x is the expression value for a given cluster, μ is the mean expression across all clusters for that gene, and σ is the corresponding standard deviation. This row-scaling approach enables direct comparison across genes with different absolute expression magnitudes. In the multi-dataset module, heatmaps can be split by dataset of origin or metadata condition, displaying average expression as side-by-side panels for direct cross-condition comparison within each cluster.

**Figure 8.**
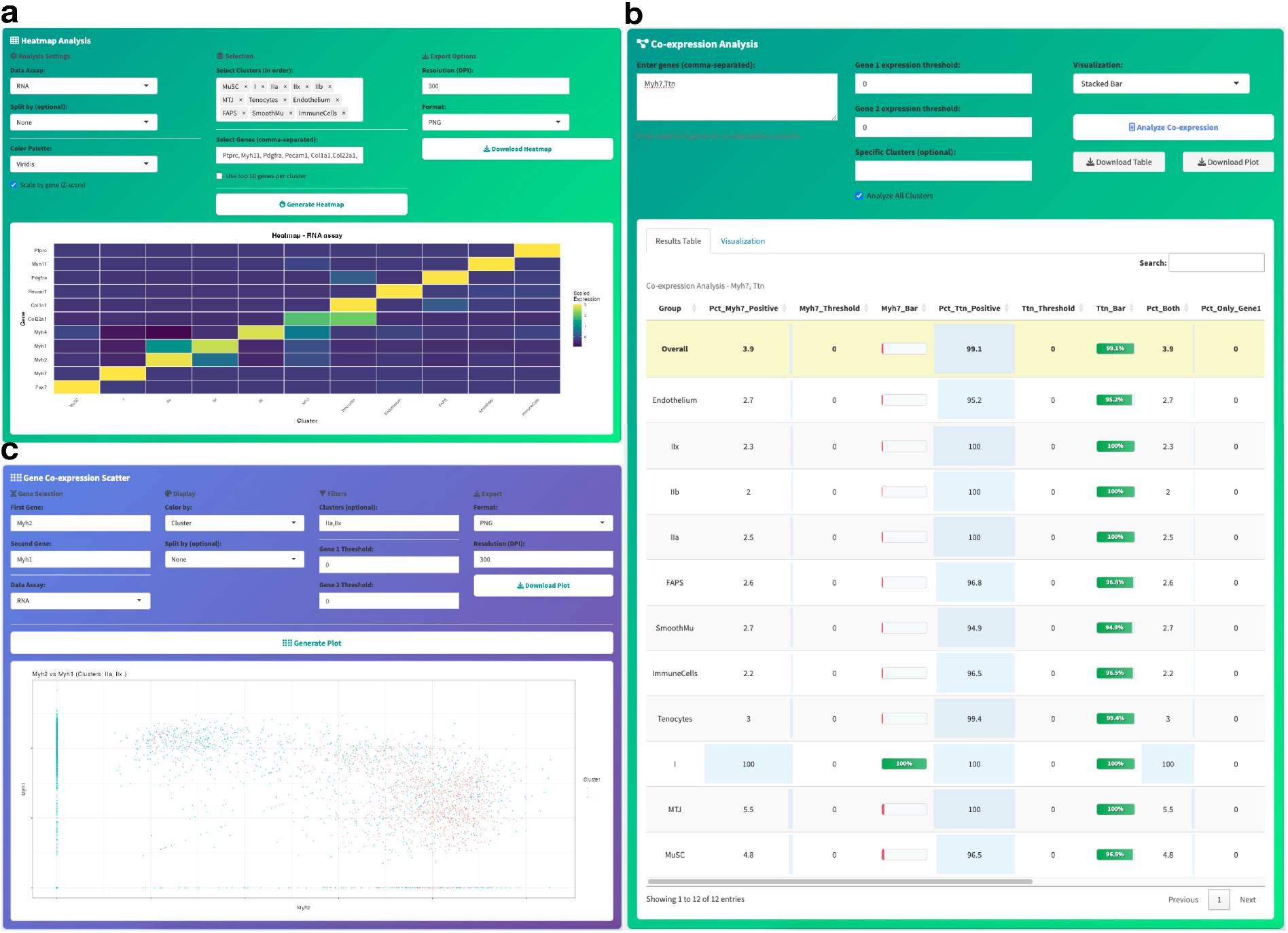
Integeated heatmap and gene co-expression analysis functionalities. (a) Heatmap analysis module enables selection of genes, clusters, assay type, and export settings, providing visualization of average expression levels across cell populations, (b) Tabular co-expression analysis summarizes expression frequency, mean values, and percentage thresholds for pairs of selected genes within defined clusters, (c) Co-expression scatter plot module depicts the relationship between two genes across clusters, with color coding according to cluster identity and support for dynamic filtering and export. These modules facilitate comprehensive exploration of combinatorial expression patterns and provide customizable export options for downstream analysis.

The co-expression analysis module identifies cells expressing multiple genes simultaneously using threshold-based binary classification (Fig. 8b). Each cell is assigned to one of four groups: both genes positive, Gene 1 only, Gene 2 only, or neither, based on user-defined expression thresholds (default: normalized expression > 0, adjustable to filter low-level background expression). In the representative example, Ttn and Myh7 display high co- expression in a subset of nuclei, illustrating the module’s ability to identify double-positive populations within a cluster.

The scatter plot module (Fig. 8c) provides single-cell resolution visualization of gene-pair relationships, with each point representing one cell colored by cluster identity. In the example shown, Myh1 and Myh2 display largely non-overlapping expression patterns across clusters, demonstrating the module’s utility for visualizing mutually exclusive gene relationships at single-cell resolution.

#### Cluster Annotation and Advanced Visualization

Following clustering and marker gene analysis, Cell-Hub enables biological interpretation through an interactive cluster annotation interface (Fig. 9a). Users assign cell type identities to computational clusters based on marker gene expression profiles, with options to customize cluster names, plot titles, label sizes, and cluster-specific colors. The annotated UMAP projection (Fig. 9b) displays assigned cell type identities directly on the embedding, facilitating interpretation of cellular composition and inter-cluster relationships.

**Figure 9.**
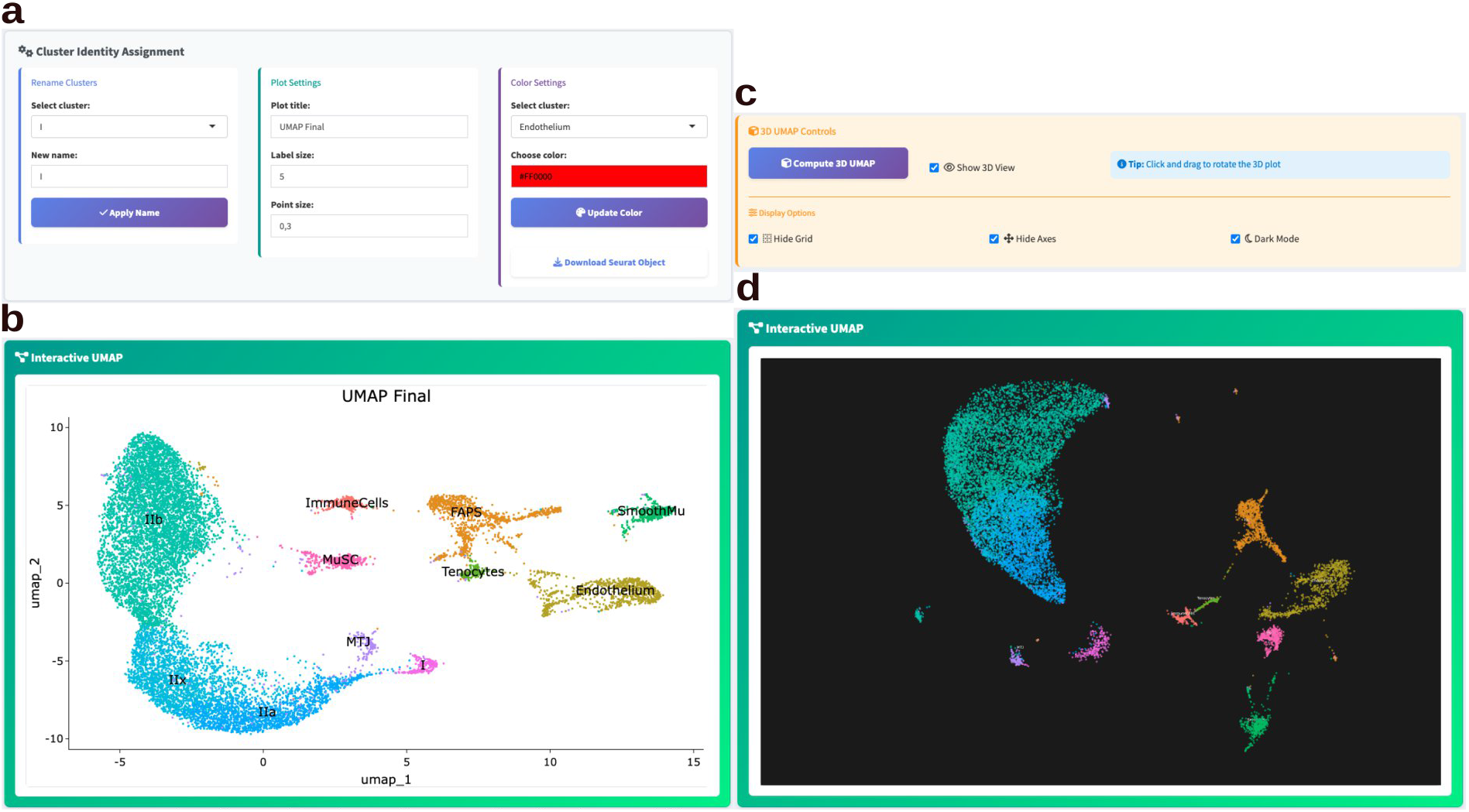
Interactive cluster identity and UMAP visualization workflow. (a) Cluster identity assignment panel allows users to rename clusters, customize plot titles, adjust label and point sizes, and assign cluster-specific colors, (b) Interactive UMAP module displays final embedding with cluster identities annotated directly on the plot, (c) 3D UMAP controls provide options to compute the three-dimensional embedding, toggle display settings, and activate dark mode for enhanced contrast, (d) 3D interactive UMAP plot visualizes cellular clusters in a dynamic, dark-mode environment for improved spatial interpretation. This workflow enables precise customization of cluster presentation and flexible, high-resolution export options for downstream analysis.

For enhanced spatial exploration, Cell-Hub implements three-dimensional UMAP visualization (Fig. 9c-d). The 3D embedding can reveal population relationships that appear compressed in two-dimensional projections, and supports interactive rotation and zoom for comprehensive examination of cluster topology. Display options include grid removal, axis hiding, and dark mode for improved contrast (Fig. 9d). All visualization settings support high-resolution export for integration into manuscripts and presentations.

#### Exclusive Biomarker Identification

Cell-Hub implements a custom exclusive biomarker detection module to identify genes with highly restricted expression in target populations. This approach is particularly valuable when standard differential expression yields numerous weakly specific genes, prioritizing markers suitable for unambiguous cell type identification or experimental validation. Unlike conventional marker identification, this module enforces strict thresholds in both target and non-target clusters to maximize specificity (Fig. 10a).

**Figure 10.**
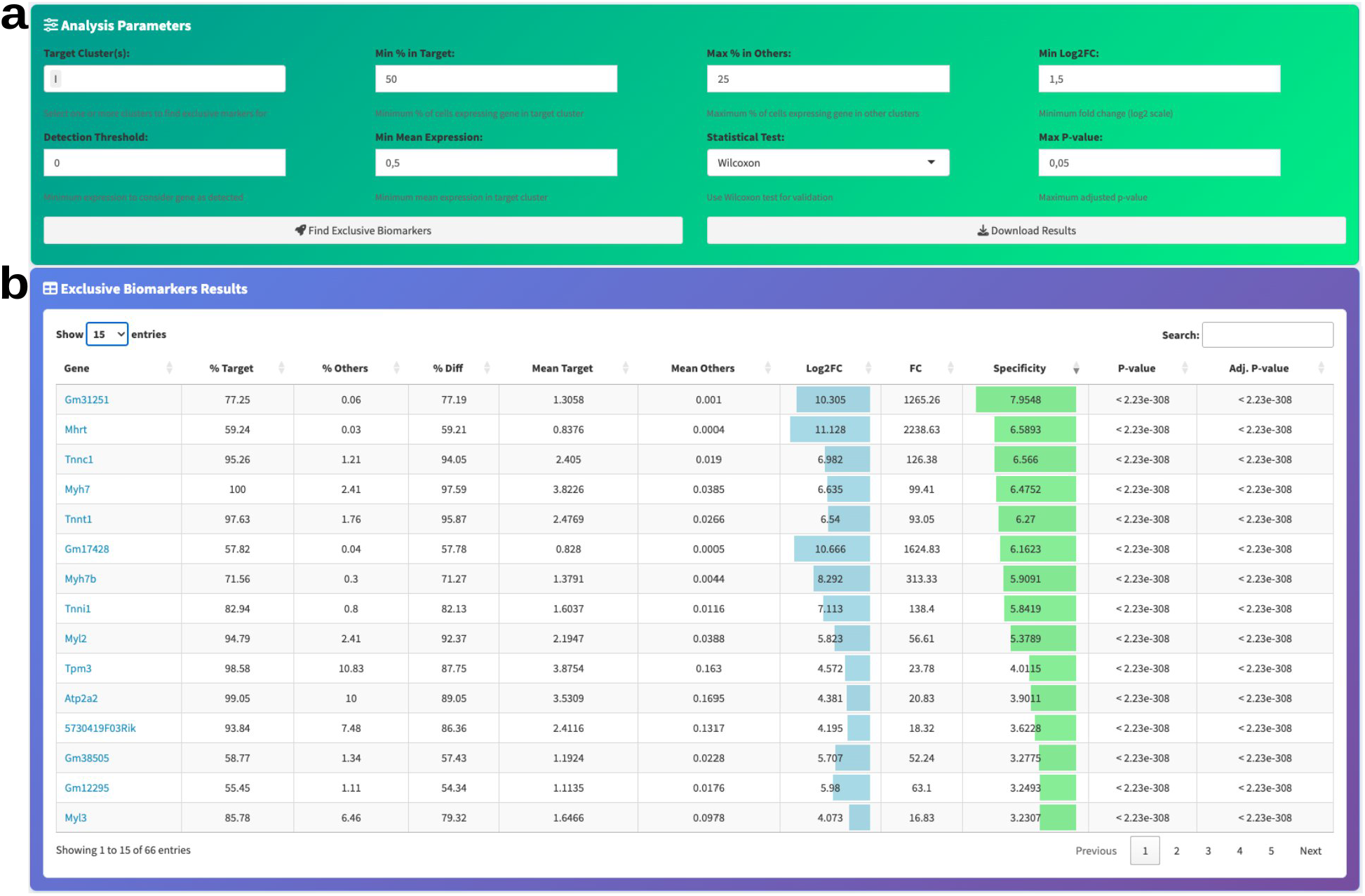
Exclusive biomarker identification. (a) Adjustable parameters for cluster-specific biomarker detection including expression thresholds, fold-change cutoffs, and statistical test selection, (b) Results table showing identified biomarkers with expression statistics, fold-change values, specificity scores, and significance metrics.

Key parameters include minimum expression percentage in target clusters (default: 50%), maximum expression percentage in other clusters (default: 25%), minimum log2 fold-change (default: 1.5), detection threshold (default: 0), and minimum mean expression level (default: 0.5 in log-normalized space). Statistical significance is assessed using the Wilcoxon rank-sum test with Benjamini-Hochberg false discovery rate correction, ensuring robust control of false positives across thousands of genes tested (user-adjustable p-value threshold, default: 0.05).

The results table (Fig. 10b) displays comprehensive metrics for each identified biomarker: expression percentages in target versus other clusters (% Target, % Others), percentage difference (% Diff), mean expression values in each group, log2 fold-change, linear fold- change, specificity score, and both raw and adjusted p-values. The specificity score quantifies biomarker exclusivity using the formula: Specificity Score = [(% Target − % Others) / 100] × log₂FC. This composite metric weights expression magnitude by detection rate differential, ranging from 0 (no specificity) to the log₂FC value (achieved when 100% of target cells and 0% of non-target cells express the gene).

In the representative example analyzing muscle populations (Fig. 10b), Myh7 (myosin heavy chain 7) shows 100% expression in target cells with only 2.41% in others (97.59% difference, specificity score = 6.48), and Tnnt1 (troponin T1) displays 97.63% versus 1.76% (95.87% difference, specificity score = 6.27), with linear fold-changes up to 2,238×, illustrating the module’s ability to resolve highly specific population markers.

#### Comparative Marker Analysis and Visualization

For comparative analysis of differential expression results, Cell-Hub provides Venn diagram comparisons (Fig. 11a) and volcano plots for detailed pairwise analysis (Fig. 11b). These visualization modules operate on previously generated differential expression results stored as tables, enabling flexible re-visualization without re-computation.

**Figure 11.**
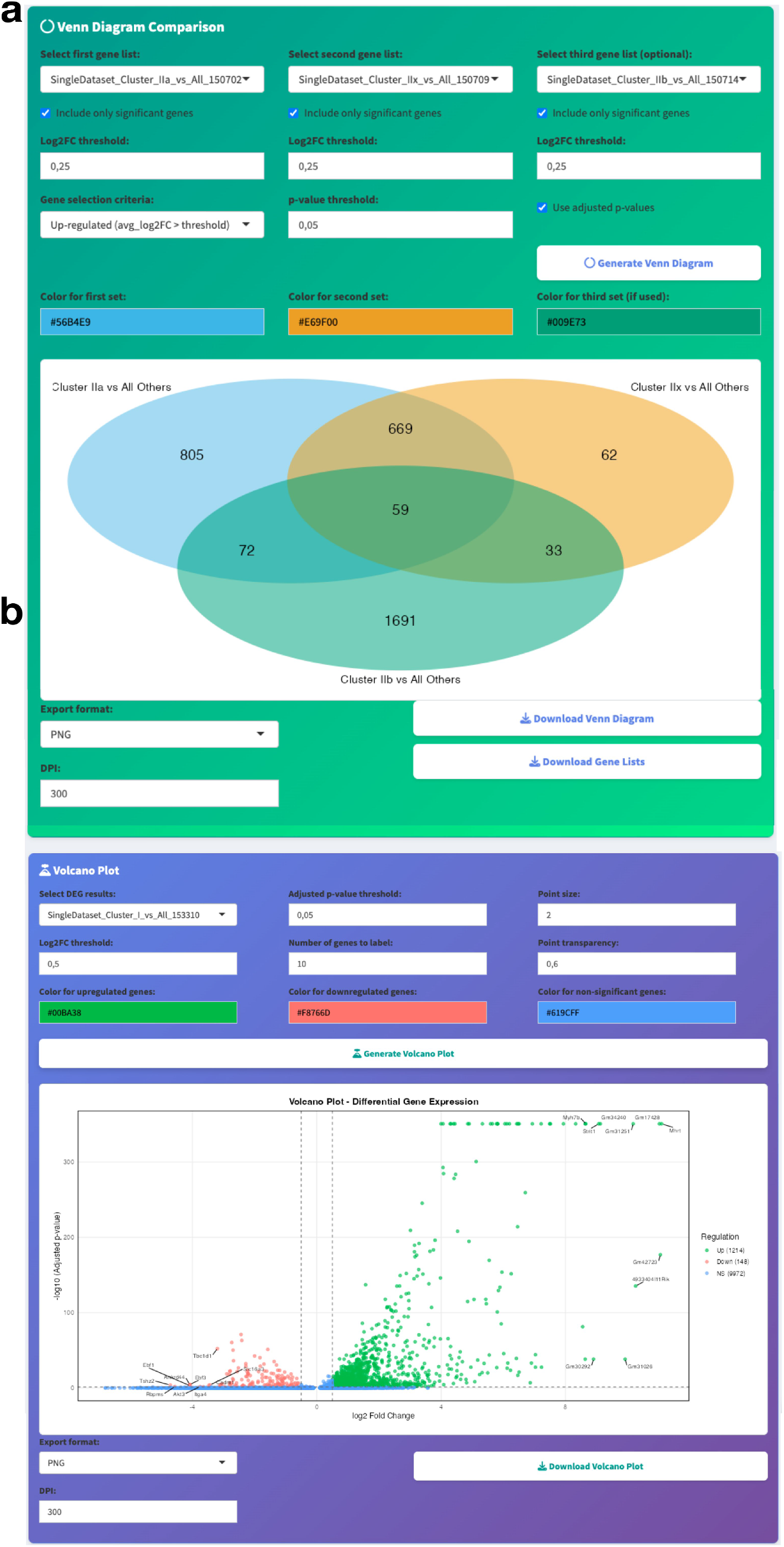
Differential gene expression visualization modules. (a) Venn diagram comparison tool enables intersection analysis of significant marker genes across multiple clusters using customizable statistical thresholds and color coding, (b) Volcano plot module provides interactive visualization of differentially expressed genes for selected clusters, with options to adjust Iog2 fold change and statistical parameters, highlight top genes, and export publication- quality plots. These modules facilitate comprehensive interpretation of transcriptomic signatures and identification of cluster-specific markers.

The Venn diagram module reveals shared versus exclusive transcriptional signatures across selected clusters. Users can generate new marker gene lists directly within the module or use previously computed results. Independent thresholds are configurable for each gene list: log2 fold-change (default: 0.5), adjusted p-value (default: 0.05, Benjamini-Hochberg correction), and directionality (upregulated/downregulated/both). Color-coded intersections display gene counts in each region, facilitating identification of pan-cluster markers (shared genes) versus population-specific markers (exclusive genes). In the representative example, 59 genes are shared across all three clusters, while 805, 62, and 1,691 genes are exclusive to each respective cluster, illustrating the module’s ability to delineate shared and population-specific transcriptional signatures simultaneously.

While Venn diagrams reveal set relationships across multiple clusters simultaneously, volcano plots enable detailed examination of individual pairwise comparisons with gene-level statistics. Each gene is positioned by log2 fold-change (x-axis) and −log10 adjusted p-value (y-axis, Benjamini-Hochberg correction), with color-coding by regulation status: upregulated (green) or downregulated (red). Users configure the p-value threshold (default: 0.05), log2FC threshold (default: 0.5), and the number of top genes to label automatically (default: 10, ranked by combined significance and fold-change magnitude).

#### Object Management: Metadata, Reductions, and Memory Optimization

In the single-dataset module, cluster identities are managed through a dedicated renaming interface that updates the active identity class and a corresponding metadata column (cluster_names) directly. A single-step undo mechanism allows reverting the most recent renaming or merging operation, providing a safety net during iterative annotation without requiring a full versioned history.

In the multi-dataset module, custom metadata fields can be defined directly through the interface and propagated across cells using any existing categorical column as a splitting variable (for example, by dataset of origin or by integration group). This enables researchers to attach experimental annotations such as condition, genotype, time point, or treatment to specific subsets of cells without external file manipulation. All metadata columns added through this mechanism are immediately available across visualization, subsetting, and statistical modules.

In parallel, all computed dimensional reductions (PCA, two-dimensional UMAP, three- dimensional UMAP, and integration-derived embeddings such as Harmony- or CCA/RPCA/MNN-corrected UMAPs) are retained within the Seurat object and exposed through a unified reduction selector. Users may switch the active embedding at any stage to compare how cell populations organize under alternative reduction strategies, without recomputation.

To mitigate the memory cost of accumulating intermediate analytical layers, Cell-Hub provides a memory optimization step in both the single-dataset and multi-dataset modules that selectively drops redundant assays (alternative assays produced by SCTransform or integration) and the scale.data layer, while preserving the RNA assay, raw counts, all dimensional reductions, neighborhood graphs, and the complete metadata. This optimization is offered as a one-click option after data loading, integration, and principal component analysis, and is recommended by default. It enables interactive analysis of larger datasets on standard workstation hardware without compromising downstream analyses such as differential expression or cell-cell communication inference.

#### Interactive Cell Subsetting and Data Export

Cell-Hub offers data subsetting for focused downstream analysis of specific cell populations or expression-defined groups. The subsetting workflow begins with visualization of the complete annotated dataset (Fig. 12a), providing context for selection decisions. Two complementary subsetting strategies are available: cluster-based extraction (Fig. 12b) allows selection of one or multiple annotated clusters by identity, while expression-based subsetting (Fig. 12c) enables threshold-driven extraction of cells expressing specific gene combinations. For expression-based subsetting, users define an expression threshold (default: 1 in log- normalized space), specify target genes as a comma-separated list, and set the minimum number of genes that must exceed the threshold per cell.

**Figure 12.**
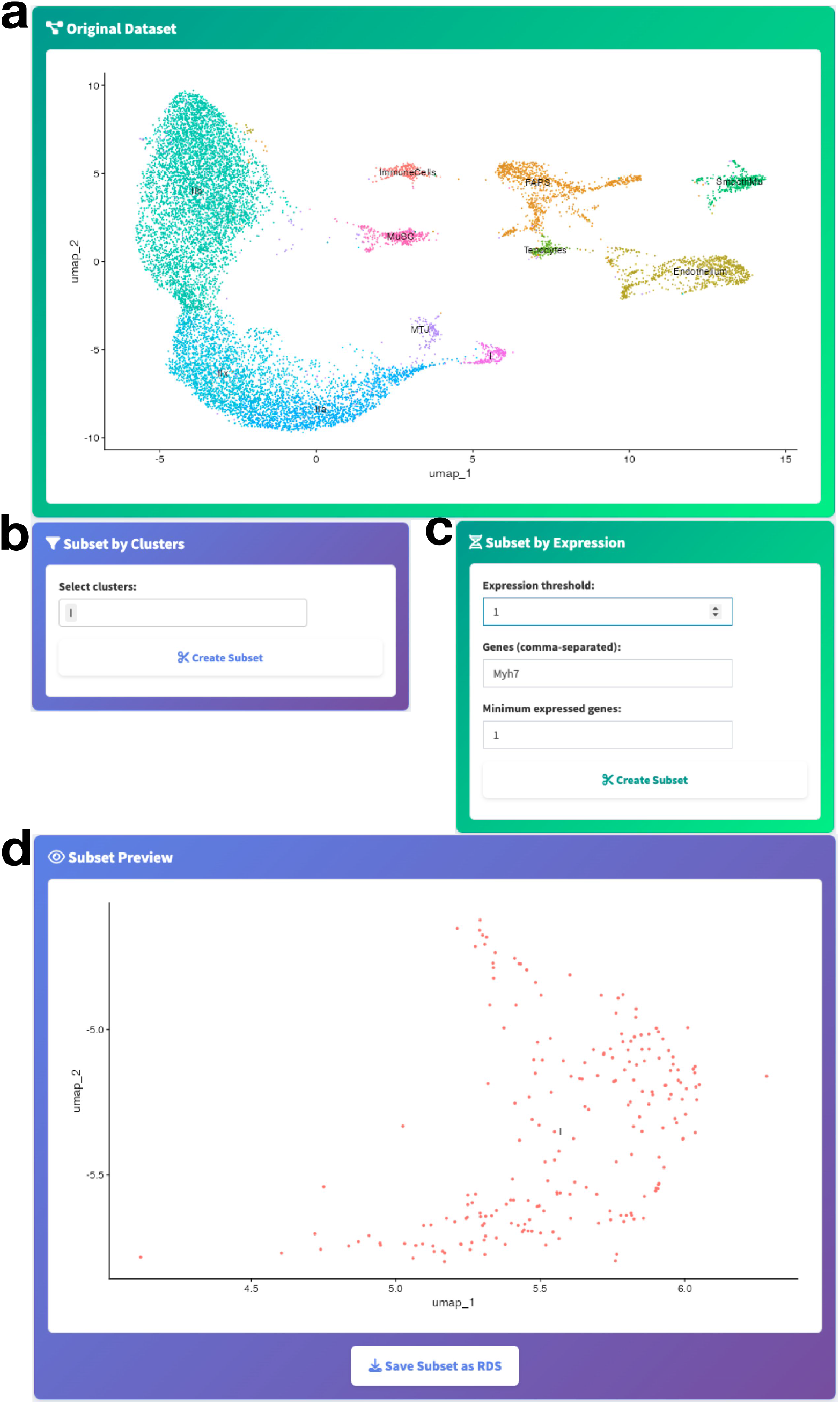
Cell Subsetting Panels. (a) UMAP visualization of the original dataset showing cluster annotations. (b) Cluster selection panel for extracting specific clusters of interest, (c) Expression-based subsetting panel allowing threshold-based extraction of cells by gene expression, (d) Preview panel displaying the UMAP projection of the selected subset with option to export.

The subset preview panel (Fig. 12d) displays the UMAP projection of selected cells, enabling visual confirmation of coherence and completeness before export. Subsets are saved as standalone Seurat RDS objects, preserving all metadata, dimensional reductions, and normalized expression matrices for independent re-analysis, integration with other datasets, or sharing with collaborators. This functionality supports iterative refinement of cell type definitions, focused differential expression on specific populations, and extraction of targeted cell subsets for specialized analyses such as trajectory inference or ligand-receptor modeling.

#### Multi-Dataset Integration and Comparative Analysis

In addition to single-dataset analysis, Cell-Hub supports the simultaneous integration of multiple scRNA-seq/snRNA-seq datasets within a unified workspace. Three integration strategies are available: CCA-based anchor integration^2^ for strong batch correction, Harmony^12^ for computationally efficient correction of technical variation, and direct merging without batch correction for datasets generated under identical experimental conditions (Fig. 13a-d). Pre-integrated Seurat objects can also be imported directly.

**Figure 13.**
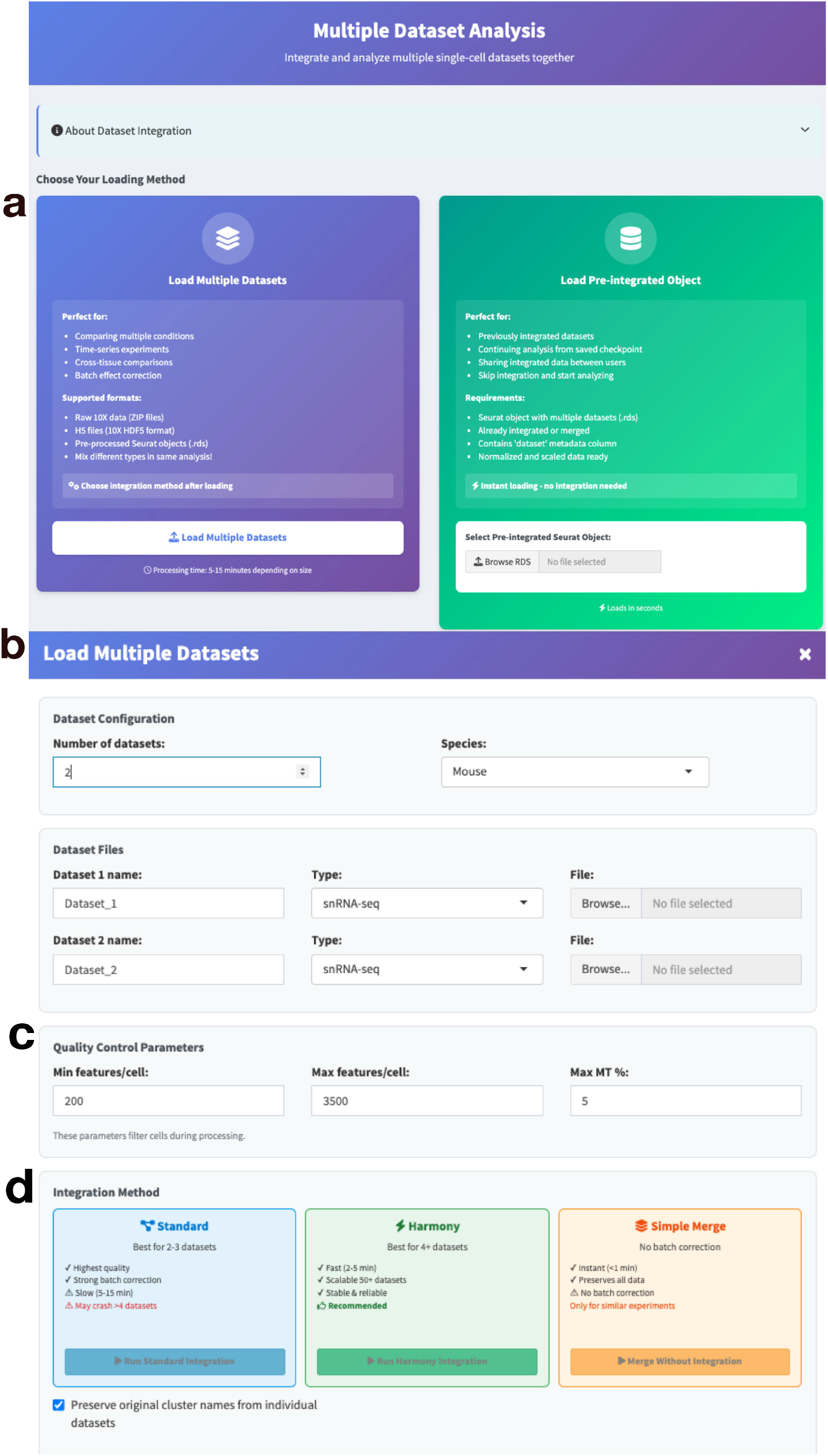
Multiple dataset integration workflow. (a) Integration method selection panel, offering choice between loading multiple raw datasets for integration or loading a pre-integrated object, (b) Dataset configuration dialog specifying number of datasets, species, dataset names, data types, and file uploads, (c) Quality control parameter input for filtering cells based on feature count, feature-to-cell ratio, and mitochondrial gene expression, (d) Integration method selection with options for Standard (batch correction), Harmony (scalable integration), and Simple Merge, with recommendations based on dataset characteristics.

A central feature of the multi-dataset module is flexible metadata management: users assign group labels to map any experimental variable (condition, genotype, time point, or treatment) onto cells across datasets. This grouping system drives all downstream comparative analyses and can be updated at any stage of the workflow.

All visualization types, including feature plots, violin plots, dot plots, ridge plots, and heatmaps, can be stratified by any metadata variable, enabling direct side-by-side comparison of gene expression patterns across the full experimental design within a single figure.

Comparative differential expression is supported through four frameworks: cluster- versus-all within the integrated object, group-versus-group contrasting any two user- defined conditions, metadata-based comparison across any combination of dataset origin or custom annotation, and within-cluster cross-condition comparison to isolate condition effects within a defined cell population. Cell population composition analysis further quantifies shifts in cluster proportions across conditions, displayed as stacked bar charts or pie charts.

#### Cell-Cell Communication Analysis with GaspouDB Integration

Cell-Hub integrates CellChat^4^ to infer intercellular communication networks from scRNA-seq data by identifying co-expressed ligand-receptor pairs across cell type dyads. Users upload previously analyzed Seurat objects containing normalized expression data and cell type annotations (Fig. 14a).

**Figure 14.**
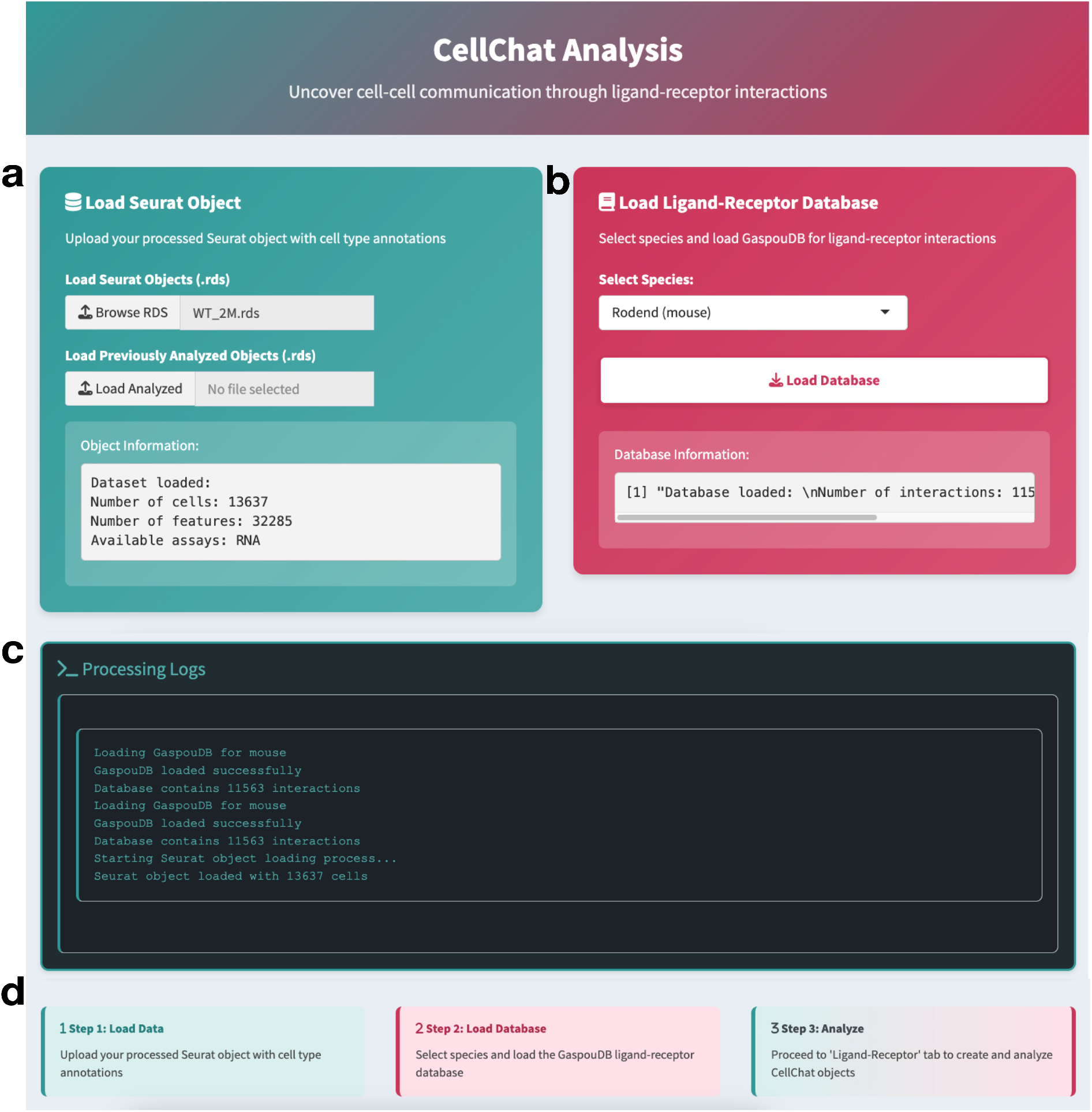
CeHChat initialization panels. (a) Data loading section for importing processed Seurat objects with cell type annotations and displaying dataset summary information, (b) Ligand-receptor database panel for selecting the species and loading interaction data from GaspouDB, with summary of available interactions, (c) Processing log section displaying status messages from data and database loading steps, (d) Stepwise instructions guiding the user through data upload, database selection, and progression to cell-cell communication analysis.

A major innovation of Cell-Hub is GaspouDB, a comprehensive ligand-receptor interaction database consolidating curated interactions from CellChat^4^, CellPhoneDB^5^, CellTalkDB^6^, and MultiNicheNet^7^ (Fig. 14b). GaspouDB contains 11,563 mouse interactions and 9,604 human interactions, substantially expanding coverage compared to any individual source database. Users select the target species, triggering automatic loading of the corresponding species- specific interactions. Processing logs confirm successful database initialization and report key metrics on the loaded interaction set (Fig. 14c-d).

#### CellChat Object Creation and Communication Inference

Following database initialization, Cell-Hub creates CellChat objects for communication network inference (Fig. 15a). Users configure the cell grouping strategy (by cluster identity or any metadata variable), apply optional filtering to include specific clusters or experimental conditions, and optionally combine multiple metadata columns into a composite grouping variable for comparative analyses.

**Figure 15.**
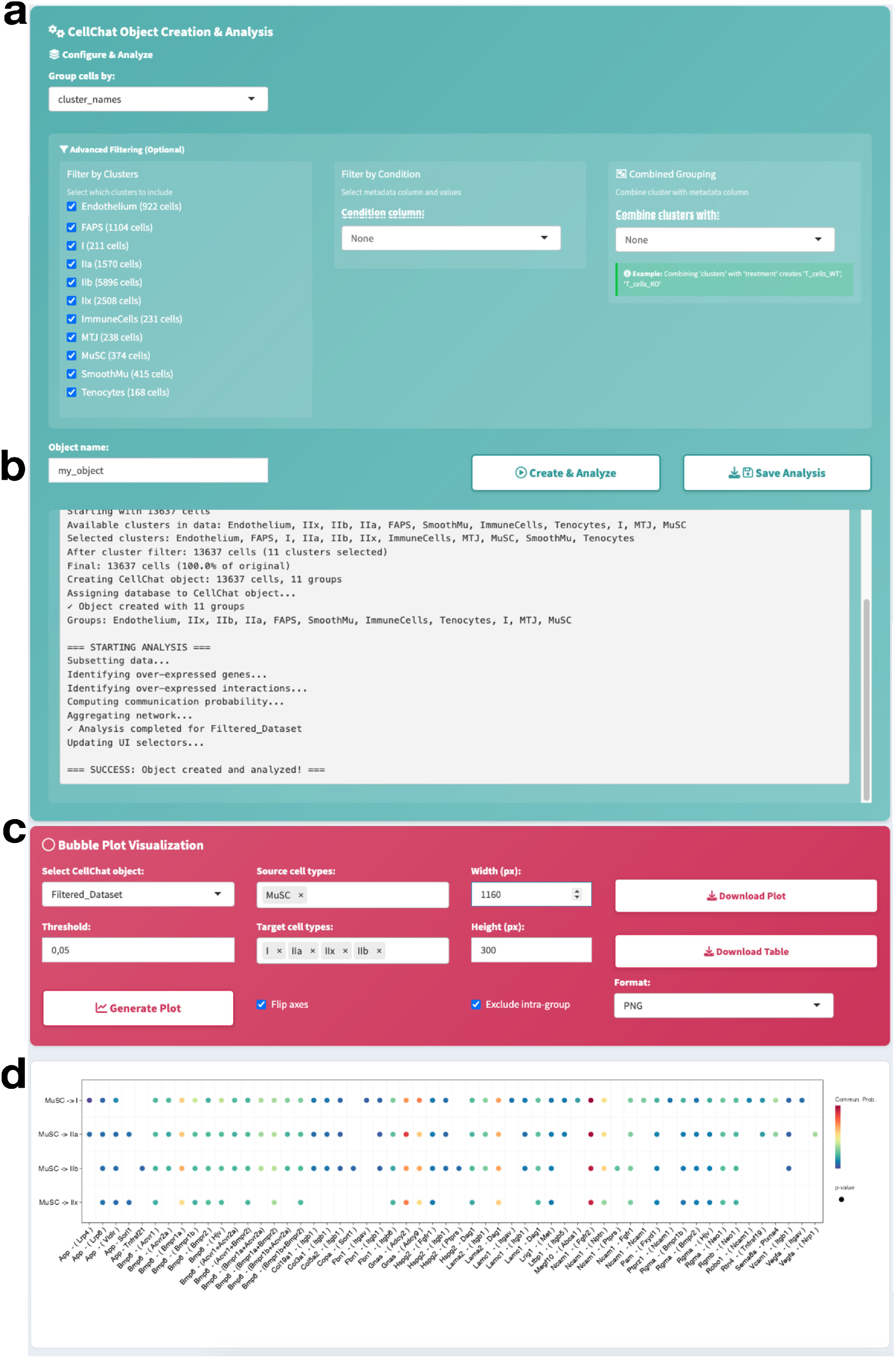
CellChat object creation and ligand-receptor analysis panels. (a) Configuration panel for grouping, filtering by clusters and conditions, and optionally combining metadata columns for analysis, (b) Analysis log showing selection of clusters, filtering steps, and status of communication network computation, (c) Bubble plot setup panel for selecting CellChat object, source and target cell types, and visualization settings, (d) Bubble plot output visualizing ligand-receptor interactions between muscle stem cells and target clusters, with dot size corresponding to communication probability.

The analysis log (Fig. 15b) provides transparent feedback on computation steps: cluster selection, database assignment, identification of overexpressed ligand-receptor pairs, computation of communication probabilities based on co-expression levels, and network aggregation. The log confirms successful object creation, enabling users to verify analysis parameters before visualization.

Cell-cell communication patterns are visualized through bubble plots (Fig. 15c-d), displaying predicted ligand-receptor interactions between source and target populations. Users select source cell types (e.g., MuSC, Muscle Stem Cells) and target cell types (e.g., fiber subtypes I, IIa, IIb, IIx), and configure a probability threshold (default: 0.05) to filter low-confidence interactions and plot dimensions. The resulting bubble plot (Fig. 15d) displays each source- target pair as a row and each ligand-receptor interaction as a column, with dot size representing communication probability and color indicating relative strength, illustrating the module’s ability to resolve distinct interaction profiles across multiple target populations simultaneously.

#### Network Visualization with Chord Diagrams

Cell-Hub provides chord diagram visualization to display intercellular communication networks predicted by CellChat (Fig. 16). Users configure sender and receiver cell types, a communication probability threshold to filter weak interactions (default: 0.05), and colors for each population (Fig. 16a).

**Figure 16.**
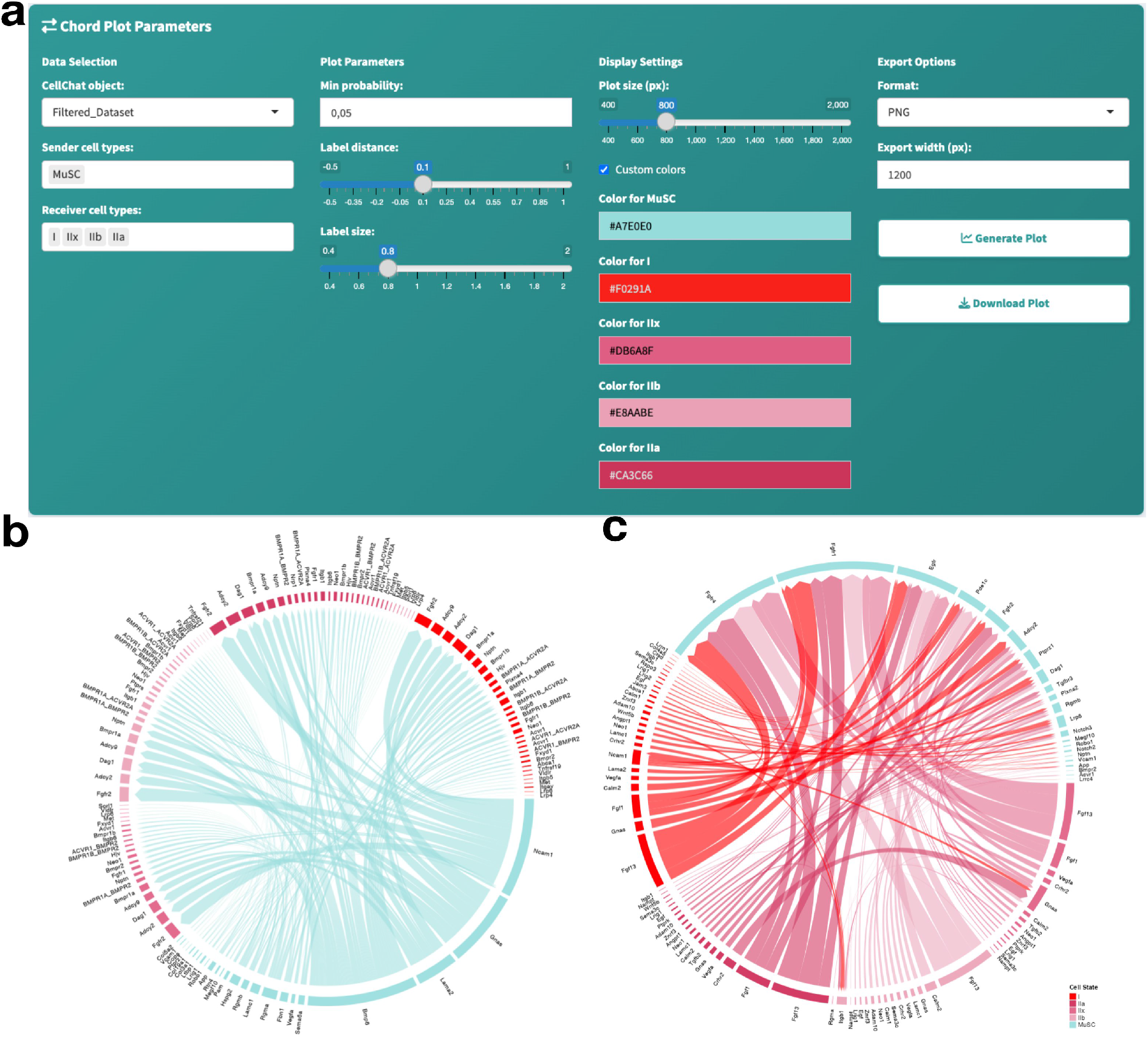
Chord plot generation and visualization panels. (a) Chord plot parameter selection panel, including CellChat object, sender and receiver cell types, probability threshold, label settings, color customization, and export options, (b) Chord plot displaying all predicted ligand-receptor interactions among selected cell types, with interaction links colored by sender and receiver groups, (c) Chord plot highlighting interactions sourced specifically from population I, using custom coloration for improved interpretability.

Chord diagrams represent cell types as segments along a circular layout, with connecting ribbons indicating predicted ligand-receptor interactions (Fig. 16b). Ribbon width represents communication probability calculated from ligand and receptor co-expression levels. Ribbon colors distinguish sender populations, revealing overall connectivity patterns across multiple cell types simultaneously.

Users can highlight outgoing interactions from specific source populations (Fig. 16c), displaying selected connections in color while dimming others. In the representative example, muscle stem cells (MuSC) display extensive predicted signaling toward multiple muscle fiber subtypes, illustrating the module’s ability to identify dominant sender populations within a communication network. Interactions are predicted based on co-expression patterns; experimental validation is required for functional confirmation.

#### Trajectory Inference with Monocle 3

Cell-Hub integrates Monocle 3^3^ to reconstruct developmental trajectories and infer pseudotemporal ordering of cells along differentiation paths. The workflow begins with conversion of analyzed Seurat objects to Monocle 3 format (Fig. 17a), preserving dimensional reductions, cluster assignments, and normalized expression data.

**Figure 17.**
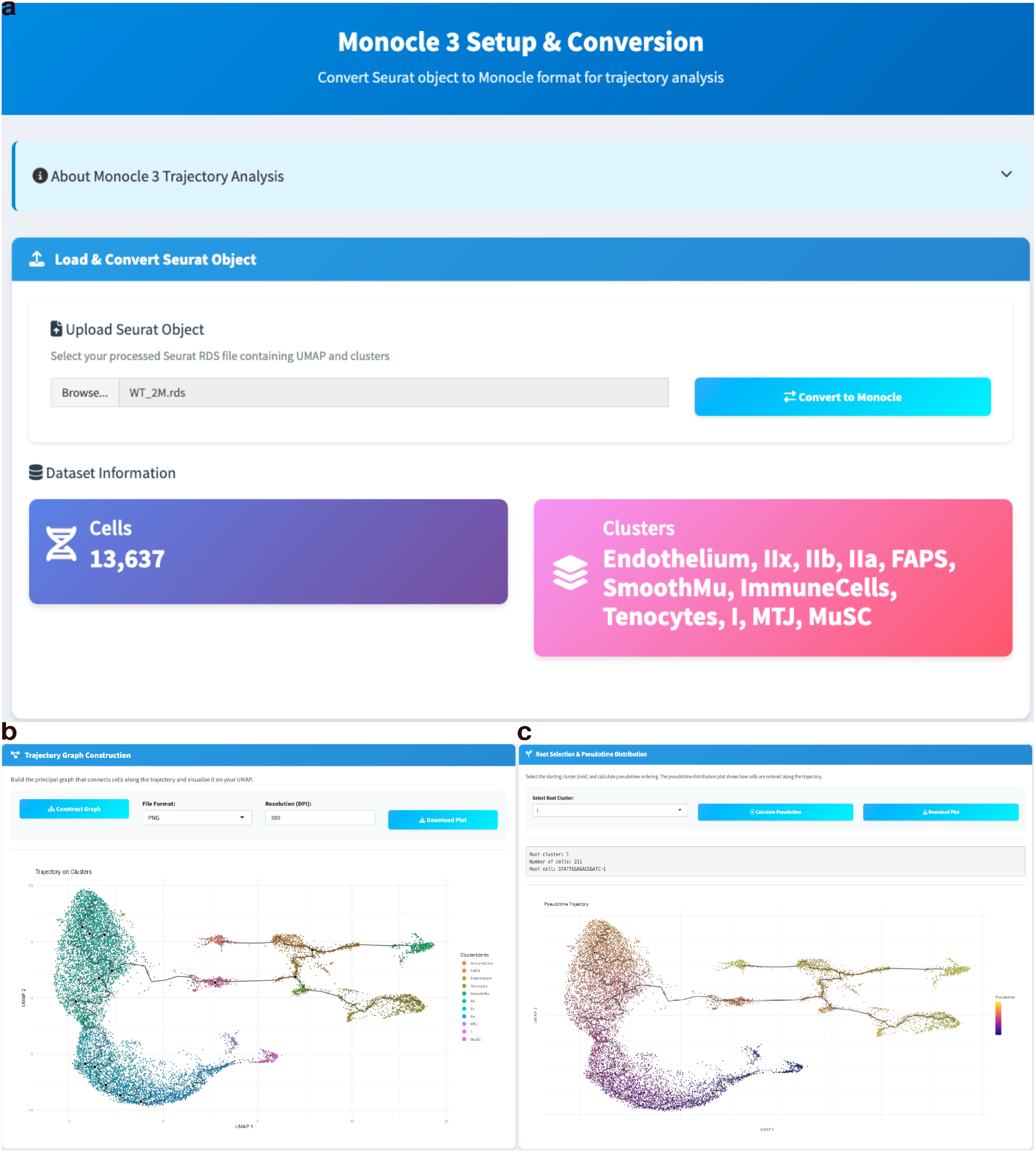
Monocle 3 trajectory analysis panels. (a) Seurat-to-Monocle conversion panel showing loaded dataset information including cell count and identified clusters, (b) Trajectory graph construction tab with options to build principal graph from cell clusters and export visualization in selected format and resolution, (c) Root cluster and pseudotime distribution panel for selecting starting point of trajectory and visualizing pseudotime progression with color gradient overlay.

The trajectory graph construction module (Fig. 17b) builds a principal graph connecting cells through their transcriptional neighborhoods in reduced-dimensional space, displayed as branching structures overlaid on the UMAP projection. Each branch represents a potential lineage trajectory, with branch points indicating divergence toward distinct terminal states. Users select a root cluster representing the earliest developmental stage based on prior biological knowledge or progenitor marker expression (Fig. 17d). Pseudotime values are then calculated from this root, representing transcriptional progression along the trajectory rather than chronological time. Cells are visualized with a continuous color gradient from early (purple) to late (yellow) pseudotime values (Fig. 17c). In the representative example, the trajectory displays branching from a progenitor population toward multiple terminal states, illustrating the module’s ability to resolve lineage relationships and order cells along differentiation paths.

#### Gene Expression Dynamics Along Trajectories

Following trajectory reconstruction, Cell-Hub identifies genes with dynamic expression patterns along pseudotime (Fig. 18). The pseudotime correlation analysis module (Fig. 18a) tests each gene for significant association with pseudotime using graph-based autocorrelation (Moran’s I statistic), reporting correlation coefficients, p-values, and q-values for multiple testing correction. Genes with significant pseudotime association are ranked for further investigation of temporal expression dynamics.

**Figure 18.**
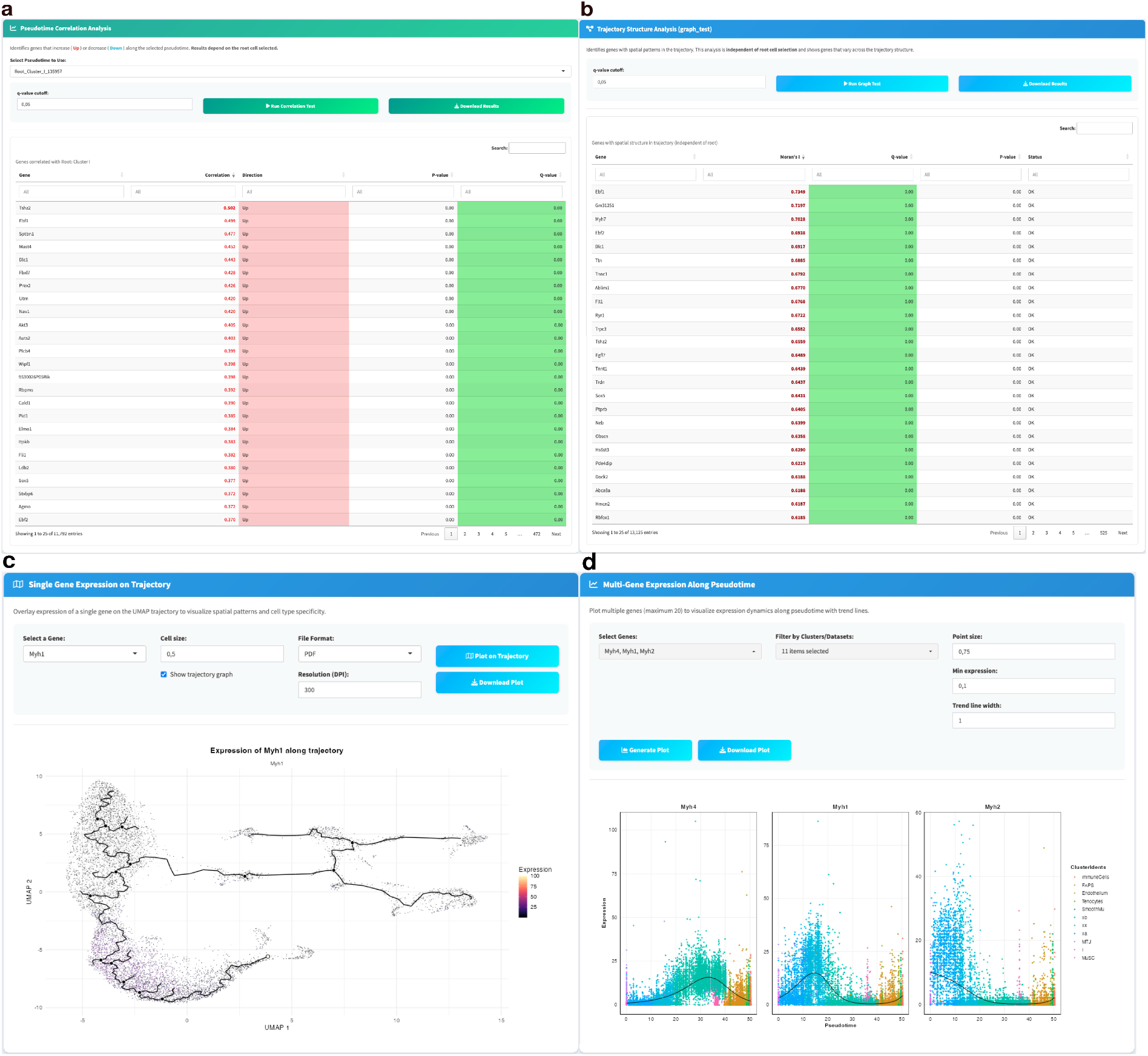
Trajectory analysis and gene expression panels. (a) Pseudotime correlation analysis table showing gene statistics and correlation patterns across the trajectory, (b) Trajectory-specific graph construction results table with gene significance metrics and correlation values, (c) Single gene expression visualization on trajectory, displaying spatial expression patterns of selected genes along the pseudotime progression with color intensity reflecting expression level, (d) Multi-gene expression along pseudotime panel showing time-series patterns for multiple selected genes with histograms representing gene expression distribution across trajectory segments and cluster composition.

The trajectory structure analysis module (Fig. 18b) identifies branch-dependent genes that distinguish alternative lineages, ranking genes by Moran’s I statistic and q-values to quantify expression divergence across trajectory branches.

Gene expression can be visualized as overlays on the trajectory graph (Fig. 18c), with a continuous color gradient representing expression intensity. In the representative example, Myh1 displays a progressive expression gradient along the trajectory, illustrating the module’s ability to resolve gene activation patterns across pseudotime. For comparative analysis across multiple genes, individual expression plots (Fig. 18d) display pseudotime on the x-axis versus expression level on the y-axis, with cells colored by cluster identity. The representative genes Myh4, Myh1, and Myh2 display distinct expression profiles across branches, demonstrating the module’s utility for distinguishing lineage-specific transcriptional programs.

#### Comparison with existing scRNA-seq GUI tools

To position Cell-Hub within the existing landscape of graphical scRNA-seq analysis platforms, we compared its features against nine widely used open-source and commercial tools (Table 1).

**Table 1:** Comparison tab feature-by-feature of Cell-Hub with nine existing GUI-based scRNA-seq analysis platforms across the major steps of single-cell analysis.

| Feature / Module | Cell-Hub ★ | ezSingleCell | OmniCellX | Asc-Seurat | ASAP | NASQAR | SCTK 2.0 | ICARUS | Cellar | SCiAp |
| --- | --- | --- | --- | --- | --- | --- | --- | --- | --- | --- |
| *Open-source tools* |  |  |  |  |  |  |  |  |  |  |
| **scRNA-seq — Preprocessing & clustering** |  |  |  |  |  |  |  |  |  |  |
| Input: 10x Cell Ranger output and H5 files | ✓ | ✓ | ✓ | ✓ | ✓ | ✓ | ✓ | ✓ | ✓ | ✓ |
| Input: RDS / Seurat object | ✓ | ✓ | — | ✓ | — | — | ✓ | — | ✓ | — |
| QC (nGenes, % mito, nCounts) and Log-normalization | ✓ | ✓ | ✓ | ✓ | ✓ | ✓ | ✓ | ✓ | ✓ | ✓ |
| SCTransform normalization | ✓ | ✓ | — | ✓ | — | — | ✓ | ✓ | — | — |
| Highly variable gene selection | ✓ | ✓ | ✓ | ✓ | ✓ | ✓ | ✓ | ✓ | ✓ | ✓ |
| PCA, UMAP | ✓ | ✓ | ✓ | ✓ | ✓ | ✓ | ✓ | ✓ | ✓ | ✓ |
| UMAP in 3D | ✓ | — | — | — | — | — | — | ✓ | — | — |
| Clustering (Louvain / SLM) | ✓ | ✓ | ✓ | ✓ | ✓ | ✓ | ✓ | ✓ | ✓ | ✓ |
| Resolution tuning / clustree | — | ✓ | — | — | — | — | — | — | — | — |
| Doublet detection with DoubletFinder | ✓ | — | — | — | — | — | ✓ | ✓ | — | — |
| Add / import custom metadata to dataset | ✓ | — | — | — | — | — | ✓ | — | ✓ | — |
| scRNA-seq — Downstream analyses |  |  |  |  |  |  |  |  |  |  |
| Exclusive biomarker discovery | ✓ | — | — | — | — | — | — | — | — | — |
| Cluster marker genes — cluster vs. rest | ✓ | ✓ | ✓ | ✓ | ✓ | ✓ | ✓ | ✓ | ✓ | ✓ |
| Pairwise DEG (cluster A vs. cluster B) | ✓ | ✓ | ✓ | ✓ | ✓ | ✓ | ✓ | ✓ | ✓ | — |
| DEG: condition vs. condition (same cluster) | ✓ | — | ✓ | — | — | — | ✓ | — | — | — |
| DEG: multi-dataset / multi-group | ✓ | — | — | — | — | — | ✓ | — | — | — |
| All pairwise cluster combinations | ✓ | — | — | — | — | — | — | — | — | — |
| Volcano plot | ✓ | ✓ | ✓ | ✓ | — | — | ✓ | — | — | — |
| Venn diagram (DEG overlap across groups) | ✓ | — | — | — | — | — | — | — | — | — |
| GSEA / pathway enrichment | under development | ✓ | ✓ | ✓ | ✓ | — | ✓ | ✓ | ✓ | ✓ |
| Heatmap (genes × clusters) | ✓ | ✓ | — | ✓ | ✓ | — | ✓ | ✓ | — | — |
| DotPlot / VlnPlot / FeaturePlot | ✓ | ✓ | ✓ | ✓ | ✓ | ✓ (no dot plot) | ✓ | — | ✓ | ✓ |
| Interactive visualization | ✓ | ✓ | ✓ | — | ✓ | — | ✓ | ✓ | ✓ | ✓ |
| Publication-ready figure export | ✓ | ✓ | ✓ | ✓ | ✓ | ✓ | ✓ | ✓ | — | ✓ |
| Cluster composition & proportions |  |  |  |  |  |  |  |  |  |  |
| Cell proportion per sample within each cluster (stacked bar) | ✓ | — | — | — | — | — | ✓ | — | — | — |
| Cell proportion per cluster within each sample | ✓ | — | — | — | — | — | ✓ | — | — | — |
| Dataset / condition origin per cluster (split UMAP) | ✓ | ✓ | — | — | — | — | ✓ | ✓ | ✓ | — |
| Cell–cell communication (CCC) |  |  |  |  |  |  |  |  |  |  |
| CCC inference tool | CellChat 2 | CellPhoneDB | CellPhoneDB | — | — | — | — | CellChat | — | — |
| Custom / proprietary LR database | GaspouDB | — | — | — | — | — | — | — | — | — |
| User-selectable LR database | ✓ | ✓ | — | — | — | — | — | — | — | — |
| Multi-condition CCC comparison | ✓ | — | — | — | — | — | — | — | — | — |
| MultiNicheNet DB (fused in GaspouDB) | ✓ | — | — | — | — | — | — | — | — | — |
| CellPhoneDB interactions in LR DB | ✓ | ✓ | ✓ | — | — | — | — | — | — | — |
| CellTalkDB interactions in LR DB | ✓ | ✓ | — | — | — | — | — | — | — | — |
| Trajectory & pseudotime |  |  |  |  |  |  |  |  |  |  |
| Pseudotime / trajectory inference | ✓ | — | ✓ | ✓ | — | — | ✓ | ✓ | — | ✓ |
| Tool used | Monocle 3 | — | PAGA | Dynverse + Slingshot | — | — | TSCAN | Monocle 3 | — | Monocle 3 |
| Pseudotime visualization | ✓ | — | ✓ | ✓ | — | — | ✓ | ✓ | — | ✓ |
| Multi-sample integration |  |  |  |  |  |  |  |  |  |  |
| Batch correction — Seurat CCA / RPCA | ✓ | ✓ | ✓ | — | — | — | ✓ | only 2 | only 2 | — |
| Batch correction — Harmony | ✓ | ✓ | ✓ | ✓ | — | — | ✓ | only 2 | ✓ | — |
| Simple merge (no batch correction) | ✓ | ✓ | ? | — | — | — | ✓ | — | — | — |
| Batch correction — fastMNN | — | ✓ | — | — | — | — | ✓ | — | — | — |
| Spatial transcriptomics |  |  |  |  |  |  |  |  |  |  |
| 10x Visium | ✓ | ✓ | — | — | — | — | — | — | ✓ | — |
| 10x Visium HD | ✓ | — | — | — | — | — | — | — | — | — |
| Spatial clustering | ✓ | ✓ | — | — | — | — | — | — | ✓ | — |

| Multi-omics & scATAC-seq |  |  |  |  |  |  |  |  |  |  |
| --- | --- | --- | --- | --- | --- | --- | --- | --- | --- | --- |
| CITE-seq (RNA + protein) | under development | ✓ | – | – | – | – | ✓ | ✓ | ✓ | – |
| 10x Multiome (RNA + ATAC) | under development | ✓ | – | – | – | – | ✓ | ✓ | – | – |
| Standalone scATAC-seq | under development | ✓ | – | – | – | – | – | – | ✓ | – |
| Deployment & accessibility |  |  |  |  |  |  |  |  |  |  |
| Docker / local install | ✓ | – | ✓ | ✓ | ✓ | ✓ | ✓ | ✓ | ✓ | – |
| Hosted web server (SaaS) | – | ? | ? | – | ? | ? | ? | ? | ? | ? |
| Output & reporting |  |  |  |  |  |  |  |  |  |  |
| Automated full analysis report (PDF / HTML) | ✓ | – | – | – | – | – | ~ | – | – | – |
| Export individual figures (PNG / SVG / PDF) | ✓ | ✓ | ✓ | ✓ | ✓ | ✓ | ✓ | ✓ | ✓ | ✓ |
| Export results tables (CSV / Excel) | ✓ | ✓ | ✓ | ✓ | ✓ | ✓ | ✓ | ✓ | – | ✓ |
| Workspace save / reload (full session) | ✓ | – | – | – | – | – | – | – | – | – |

Several of the platforms listed above are deployed as hosted web servers. Cell-Hub deliberately avoids this architecture: public instances of web-based tools are inherently constrained in the computational resources they can allocate per user, and scRNA-seq analyses routinely require tens of gigabytes of memory and substantial CPU time for integration, differential expression, cell-cell communication, and trajectory inference. Furthermore, uploading large count matrices through a browser raises data privacy concerns. Cell-Hub is therefore distributed as a Docker image that runs locally, scaling with the user’s own hardware and keeping all data on their machine.

## Discussion

Modern single-cell sequencing technologies generate datasets of unprecedented scale and complexity, creating a significant barrier for biologists without extensive computational training. While command-line tools like Seurat, Cellchat, and Monocle provide powerful analytical capabilities, their steep learning curve prevents many experimental researchers from independently analyzing their own data.

Cell-Hub addresses this accessibility challenge by providing a graphical interface that enables researchers to independently analyze data produced during their single-cell experiments. Users can clean their data, perform dimensionality reduction, identify and characterize cell types of interest, determine differentially expressed genes between conditions, explore gene expression visualizations for presentations and articles, and further advance their analysis with ligand-receptor interactions based on consolidated databases and pseudotime trajectory inference.

Several graphical interface solutions exist for single-cell analysis, with ASC-Seurat being a notable example. Each implementation incorporates the features its developers considered most relevant, but they typically cover only a subset of a complete single-cell workflow, and their interfaces are often not designed for users without prior bioinformatics experience.

In contrast, our application provides access to the most recent versions of commonly used libraries and incorporates most functionalities necessary for comprehensive single-cell analysis.

Cell-Hub’s primary innovation is the seamless integration of multiple analytical frameworks (Seurat, CellChat, Monocle) within a unified, all-inclusive workflow. Users can progress from quality control through clustering, differential expression, cell-cell communication inference, and trajectory analysis without manual data format conversion or custom integration code, a critical advantage as these tools typically operate independently and require substantial programming effort to combine. This integrated approach enables researchers to perform complete analyses from raw data to publication-ready results within a single interface, eliminating the fragmented workflows that characterize most existing solutions.

Docker-based deployment enables local installation, allowing researchers to leverage their own computational resources rather than relying on external servers or cloud infrastructure, which is particularly valuable for large datasets. Additionally, to enhance cell-cell communication analysis, Cell-Hub incorporates GaspouDB, a consolidated ligand-receptor interaction database integrating multiple curated sources.

This application represents a first step toward democratizing single-cell analysis. Ideally, a future development direction would be the packaging of Cell-Hub as a native desktop application, eliminating the need for Docker and offering improved performance and a more seamless installation experience for end users.

## Acknowledgements

We thank Edgar Jauliac and Léa Delivry for bioinformatics support, and Stéphanie Backer and Florian Britto for helpful comments on the manuscript.

## Authors’ contributions

G.M.: Conceptualization, Software, Methodology, Writing – Original Draft, Writing – Review & Editing. M.D.: Validation, Writing – Review & Editing. V.T.: Validation, Writing – Review & Editing. H.A.: Funding Acquisition, Writing – Review & Editing. P.M.: Funding Acquisition, Supervision, Writing – Review & Editing.

## Ethics, Consent to Participate, and Consent to Publish declarations

All animal experiments were conducted according to the National and European legislation and institutional guidelines for the care and use of laboratory animals approved by the French government (Ministère de l’Enseignement Supérieur et de la Recherche, autorisation APAFiS #44987-2023092714211861 v4 approved on February 5, 2024, “Evaluation of gene therapy approaches and antisense strategies for genetic diseases”). No human participants were involved in this study; therefore, consent to participate and consent to publish are not applicable.

## AI assistance declaration

During the preparation of this manuscript, the authors used Claude (Anthropic) to assist with manuscript revision and to assist in portions of the Cell-Hub software development. All content was reviewed, verified, and approved by the authors, who take full responsibility for the integrity of the work.

## Data availability statement

Cell-Hub is freely available as a Docker image at Docker Hub (gaspardmacaux/cell-hub:latest) and the source code is available at https://github.com/GaspardMacaux/Cell-Hub. GaspouDB is distributed within the Docker image and available on the GitHub repository. RNA sequencing data used for demonstration purposes have been deposited in the Gene Expression Omnibus (GEO) under accession number GSE309529 (https://www.ncbi.nlm.nih.gov/geo/query/acc.cgi?acc=GSE309529).

